# Detection of spatial chromatin accessibility patterns with inter-cellular correlations

**DOI:** 10.1101/2024.03.09.584249

**Authors:** Xiaoyang Chen, Keyi Li, Xiaoqing Wu, Zhen Li, Qun Jiang, Yanhong Wu, Rui Jiang

**Author notes:** These authors are equal contributors (Xiaoyang Chen, Keyi Li).

## Abstract

Recent advances in spatial sequencing technologies enable simultaneous capture of spatial location and chromatin accessibility of cells within intact tissue slices. Identifying peaks that display spatial variation and cellular heterogeneity is the first and key analytic task for characterizing the spatial chromatin accessibility landscape of complex tissues. Here we propose an efficient and iterative model, Descartes, for spatially variable peaks identification based on the graph of inter-cellular correlations. Through the comprehensive benchmarking for spatially variable peaks identification, we demonstrate the superiority of Descartes in revealing cellular heterogeneity and capturing tissue structure. In terms of computational efficiency, Descartes also outperforms existing methods with spatial assumptions. Utilizing the graph of inter-cellular correlations, Descartes denoises and imputes data via the neighboring relationships, enhancing the precision of downstream analysis. We further demonstrate the ability of Descartes for peak module identification by using peak-peak correlations within the graph. When applied to spatial multi-omics data, Descartes show its potential to detect gene-peak interactions, offering valuable insights into the construction of gene regulatory networks.

## Introduction

Spatial molecular profiling enables the measurement of biomolecules within intact tissue sections, facilitating the construction of spatially resolved cell atlas^1,2^, analysis of cellular communication^3,4^, and exploration of the cancer tumor microenvironment^5^. Recent innovations in spatial sequencing technologies, have integrated spatial barcoding schemes with assays for transposase-accessible chromatin using sequencing (ATAC-seq), allowing for the capture of spatial epigenetic information at the tissue level^6,7^. Moreover, spatial multi-omics sequencing enables the detection of connections between chromatin accessibility and gene expression in the spatial context, and so on, provides valuable insights into spatiotemporal gene regulatory mechanisms of complex tissues^8,9^.

A key analytic task in spatial sequencing data is to identify spatially variable (SV) features that display spatial patterns of chromatin accessibility or gene expression^10^. Numerous methods^11-19^ specifically developed for spatial RNA-seq (spRNA-seq) data have achieved notable success in identifying SV genes^20,21^, while there is still a lack of methods tailored for modeling spatial ATAC-seq data^21^ (spATAC-seq). Given the characteristic of higher sparsity in spATAC-seq data, the accessibility pattern of features (peaks in common scenarios) in a single slice is more discrete than spRNA-seq data, rendering the assumptions underlying spRNA-seq data-based methods inapplicable to spATAC-seq data. Furthermore, since the dimensionality of spATAC-seq data (the number of peaks) is an order of magnitude larger than the number of genes, methods designed for spRNA-seq data, which typically model, evaluate and rank genes individually, are notably inefficient for spATAC-seq and require even several hours for computation. On the other hand, several methods for identifying peaks with high heterogeneity, as used for single-cell ATAC-seq (scATAC-seq), such as selecting peaks with the highest degree of accessibility (commonly used in scATAC data analysis)^22-26^, a correlation-based method named Cofea^27^, and specific functions provided in analytic pipelines^28,29^, ignore spatial information and thus cannot capture the spatial variations. Intuitively, the above two types of approaches fail to take full advantage of the intrinsic information from spatial distribution and data matrices, suggesting the pressing demand for methods to identify SV peaks. Besides this, other crucial analytic tasks, such as spatially peak module identification, gene-peak interaction detection, and data imputation, also lack the tailored modeling for spATAC-seq data.

To fill these gaps, we present Descartes, a graph-based model, for DEtection of Spatial Chromatin Accessibility patteRns with inTEr-cellular correlationS. Leveraging the graph of inter-cellular correlations, Descartes adeptly evaluates and identifies SV peaks by analyzing the self-correlations of peaks within the graph. To navigate the inherent challenge of highly dispersed accessibility patterns in spATAC-seq data, Descartes incorporate chromatin accessibility information with spatial locations during graph construction, and iteratively updates the graph to capture the intricate relationships between neighboring cells. Based on comprehensive benchmarking on 16 slices from 4 datasets, we demonstrate the superiority of Descartes in identifying SV peaks that reveal cellular heterogeneity and tissue structure. Beyond its analytic advantages, Descartes also surpasses other methods with spatial assumptions in computational efficiency. By leveraging neighboring relationships of the graph, Descartes can impute data through signals from adjacent cells, thereby enhancing the accuracy of downstream analyses. Utilizing the inter-correlation of features within the graph, Descartes enables the capture of inherent relationships between features: when applied to spATAC-seq data, Descartes can obtain a peak-peak correlation matrix, facilitating peak module identification; when applied to spatial multi-omics data, Descartes can produce a gene-peak correlation matrix, enabling the detection of gene-peak interaction and facilitating the discovery of gene regulatory networks.

## Results

### The Descartes model

The Descartes model aims to identify informative peaks that simultaneously characterize cell heterogeneity at cellular level and spatial continuity at tissue level (Fig. 1). Given a peak-by-spot matrix with spatial locations of spots (also can be replaced by cells), Descartes evaluates and ranks peaks based on the graph of inter-cellular correlations, which are integrated from both spatial and chromatin accessibility information. More specifically, the procedure of Descartes can be divided in to five main steps: (i) constructing a spatial graph based on spatial locations of spots; (ii) performing principal component analysis (PCA) transformation on the peak-by-spot matrix with part of peaks to obtain the latent embeddings of spots; (iii) constructing a graph of chromatin accessibility based on the latent embeddings, and integrating with the spatial graph to obtain a graph of inter-cellular correlations; (iv) utilizing the self-correlation of each peak within the graph to calculate the importance score; (v) evaluating and ranking all peaks based on importance scores, and then feeding the current ranking back into step (ii). Descartes iteratively performs steps (ii) through (v) until the obtained ranking of peaks undergoes minimal changes. The concept of modeling employed here is akin to Moran’s I^30^, a statistical measure frequently applied to spRNA-seq data^17-19^. Descartes, however, tailors its modeling specifically to the characteristics of spATAC-seq data, integrating spatial information with chromatin accessibility data into a cohesive framework (the technical details of Descartes are provided in the “Methods” section). When applying Descartes, researchers could designate a specific number or a predetermined proportion of peaks as SV peaks, based on the ranking or importance scores of peaks. Leveraging the graph of inter-cellular correlations, Descartes can accomplish data imputation. Besides, Descartes can generate peak-peak similarity matrix based on the graph of inter-cellular correlations, and further utilize the similarity matrix for peak module identification. For spatial multi-omics data, such as the simultaneous capture of chromatin accessibility and gene expression information in a single slice, Descartes can obtain a gene-peak similarity matrix in a similar manner, enabling the detection of gene-peak interactions.

**Fig. 1.**
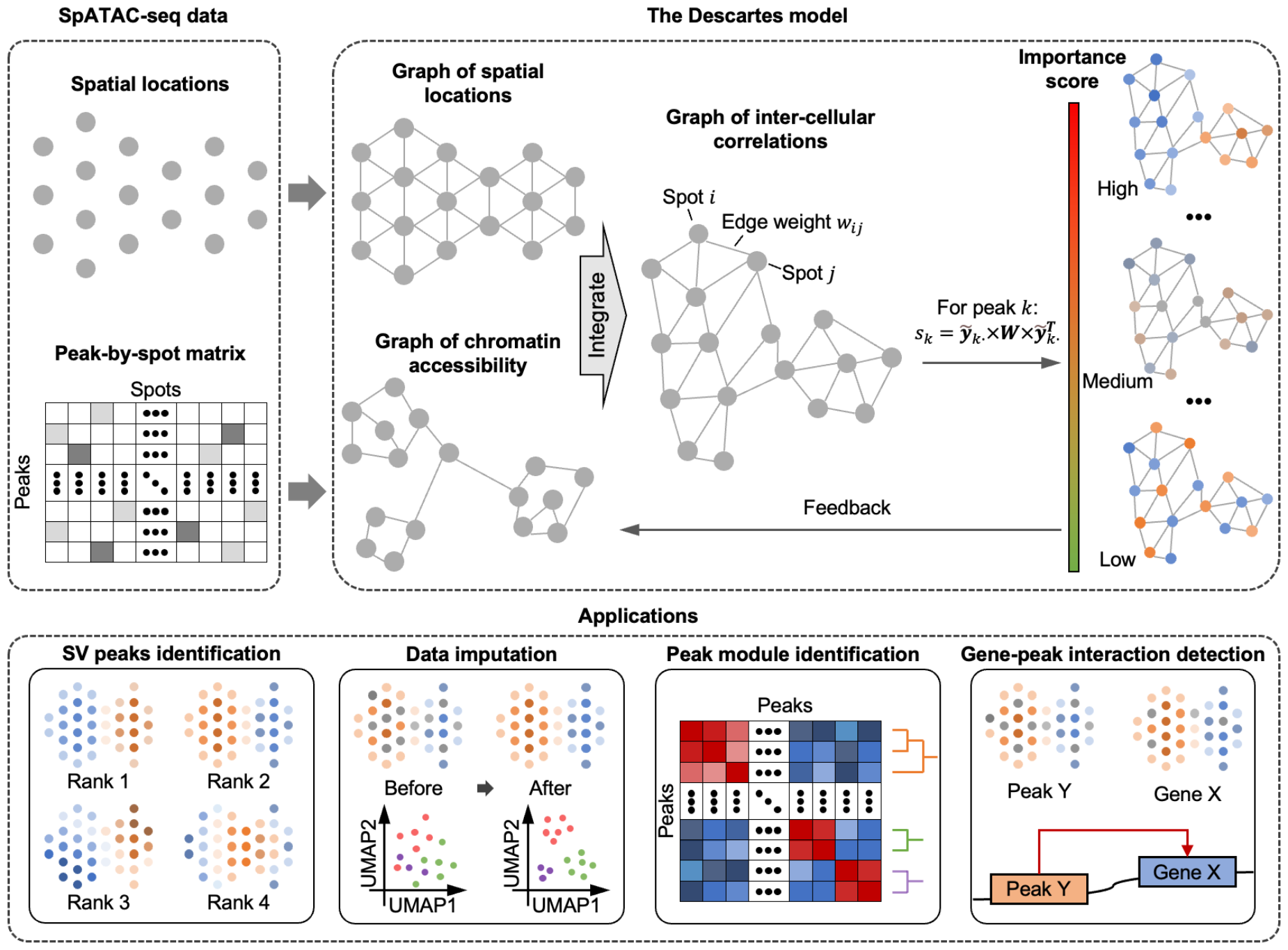
The overview of Descartes. Descartes first constructs two distinct graphs based on spatial locations of spots and the peak-by-spot matrix, that is, the graph of spatial locations and the graph of chromatin accessibility. Next, Descartes integrates the two graphs to derive the graph of inter-cellular correlations, and utilizes the self-correlation of each peak within the graph to calculate the importance score. Based on these importance scores, Descartes ranks all peaks and selects SV peaks. The SV peaks identified in each iteration are utilized to feedback and update the graph of chromatin accessibility, thereby refining the accuracy of neighborhood relationships among cells. Besides SV peaks identification, Descartes can also be applied to data imputation, peak module identification and detection of gene-peak interaction.

### Benchmarking performance of Descartes using mouse brain data

At the outset, we used the mouse brain dataset^8^, which comprises four tissue slices with well-annotated domain labels (Supplementary Table 1), to assess the performance of Descartes in SV peaks identification. Due to the absence of methods specifically designed for spATAC-seq data, Descartes was benchmarked against two types of published methods: methods tailored for spRNA-seq data, including SOMDE^15^, Moran’s I^17^, SpatialDE2^11^, SpatialDE^12^, SPARK-X^13^, SPARK^31^, scGCO^14^, Sepal^16^, and methods designed for scATAC-seq data, including directly selecting peaks with high degree of accessibility (commonly used for scATAC-seq data analysis)^22-26^, epiScanpy^28^, Signac^29^, and Cofea^27^ (Methods). Drawing from scIB^32^ and our previous work^27^, we assessed different methods from two perspectives: the ability to facilitate clustering performance and capture domain-specific signals. The evaluation process is detailed in the “Methods” section. For cell clustering performance, we employed normalized mutual information (NMI), adjusted Rand index (ARI), and adjusted mutual information (AMI) scores as metrics. To evaluate the capture of domain-specific signals, we used the overlap proportion (OP) of domain-specific peaks with SV peaks as the metric, where OP1 and OP2 correspond to overlaps identified by “tl.rank_features” function in epiScanpy and “FindAllMarkers” function in Signac, respectively. Higher scores in these metrics indicate better method performance. For a fair comparison, we tested each method by selecting 10,000 SV peaks and conducted all tests on a server with 128GB of memory and equipped with 32 units of 13th Gen Intel(R) Core(TM) i9-13900K to simulate typical personal computing conditions. SPARK and SpatialDE2 encountered memory overflow errors during the process, and scGCO did not converge even after 24 hours. Given that the mouse brain dataset represents the smallest scale within our collected data, we decided not to include these two methods in further comparisons. The benchmark results for other methods are depicted in Fig. 2a (with pre-processed results in Supplementary Fig. 1). Descartes not only gets the highest in overall scores, but also excels in all metrics of cell clustering and uncovering domain-specific signals, indicating the superiority in identifying SV peaks rich in the capture of cellular heterogeneity and tissue structure. Besides, due to lacking the incorporation of spatial information, methods based on scATAC-seq data perform significantly worse in uncovering domain-specific signals, leading to lower overall scores. In terms of computational efficiency, except for HDA, epiScanpy and Signac, which evaluate and rank peaks based on simplistic statistical information, Descartes is over two times more efficient than the other methods (Fig. 2b). Thus, Descartes demonstrates a significant advantage in both accuracy and computational efficiency in identifying SV peaks from spATAC-seq data.

**Fig. 2.**
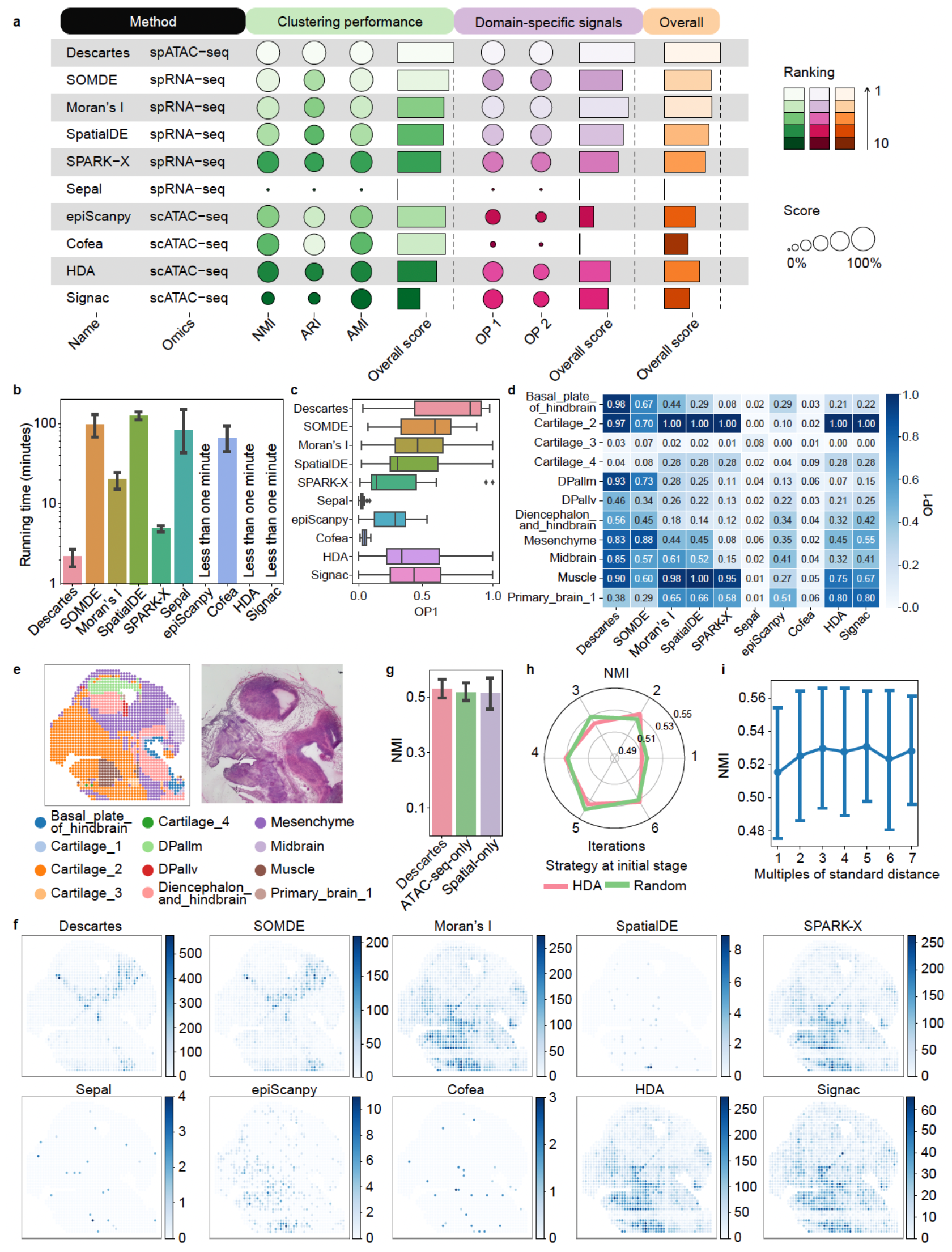
Benchmarking performance of SV peaks identification on the mouse brain dataset. **a**, Overview for benchmarking results of different methods from three perspectives, that is the ability to facilitate clustering performance, capture domain-specific signals and maintain spatial continuity (see Methods for further visualization details). **b**, Running time of different methods. **c** and **d**, Overlapped proportion of SV peaks identified by Descartes and baseline methods with domain-specific peaks related to overall domains (**c**) or each domain (**d**). Using the ‘tl.rank_features’ function in epiScanpy, we defined the top 100 peaks with the lowest p-values in each domain as domain-specific peaks. **e**, Visualization of domains within the tissue space (left) and the corresponding histological image (right). **g**, Clustering performance using SV peaks identified by Descartes and its variants. ATAC-seq-only and spatial-only represents the variants of Descartes that only utilizes the graph of chromatin accessibility and the graph of spatial locations, respectively. **h-i**, Clustering performance using SV peaks identified by Descartes with different number of iteration (**h**), different strategies for peak selection at initial stage (**h**) and different multiples of standard distance (**i**). Clustering performance is evaluated by NMI scores. In **b, g** and **i**, the error bars denote the 95% confidence interval, and the centres of the error bars denote the average value. **j**, Top-ranked SV peak identified by each method on the E13_5-S1 slice, with the raw count values visualized in the tissue space.

Next, we conducted an in-depth analysis of the benchmark results, to delve into the intrinsic differences between various methods. Taking the E13_5-S1 slice as a case study, we observed that the overall distribution of SV peaks identified by Descartes shows a higher overlap with domain-specific peaks than baseline methods (Fig. 2c). When focusing on individual domains, we found that Descartes also outperforms baseline methods in capturing cellular heterogeneity across various domains, including those with few samples, such as ‘DPallv’ (13 spots) (Fig. 2d). Domains where Descartes underperforms, such as ‘Cartilage_4’ (2 spots) and ‘Primary_brain_1’ (6 spots), typically suffered from an extremely low number of spots and the lack of spatial continuity, which may lead to excessive noise in identifying domain-specific peaks and diminish the informative value for evaluation. Similar trends could also be observed in other slices (Supplementary Fig. 2 and 3). We then visualized the top-ranked peak selected by each method, and compared it against the spatial coordinates of the domain and the corresponding histological image (Fig. 2e-f and Supplementary Fig. 4). The top-ranked peak identified by Descartes and SOMDE closely corresponds to specific tissue regions, while SOMDE, which utilizes self-organizing maps, tends to select peaks accessible over larger areas. Moran’s I and SPARK-X only capture accurate SV peaks in half of slices, and the spatial assumptions in SpatialDE and Sepal prove less applicable to spATAC-seq data. Methods based on scATAC-seq did not show clear spatial patterns in the visualization.

In our final analysis, to dissect the key to superior performances of Descartes, we conducted a series of ablation experiments and employed NMI from clustering performance as the evaluative standard. The essence of Descartes lies in the unique integration of spatial and chromatin accessibility information in graph construction. We evaluated Descartes against two variants: one utilizing only spatial information (spatial-only) and another relying only on chromatin accessibility information (ATAC-seq-only). These variants echo strategies applying in prior spRNA-seq methods^18, 19^, yet neither incorporates a fusion of these elements. As demonstrated in Fig. 2g, Descartes that considers both spatial and chromatin accessibility information achieves superior performance than any single-element-focused variant. Intriguingly, the variant ATAC-seq-only outperforms that relying only on spatial locations of spots, highlighting the discrete nature of spATAC-seq data in space and the introduction of noise when only spatially accessible pattern of individual peaks is considered. Nonetheless, the graph constructed from spatial locations is indispensable, encapsulating structural information of tissues. To mitigate noise, Descartes constructs the spatial graph using neighbors of spots within five standard deviations, akin to applying a low-pass filter to the signal in space, which proved more effective than considering only a few neighbors around a spot (Fig. 2h). For efficient and precise graph construction based on ATAC-seq matrix information, an iterative strategy was adopted in the initial graph construction and updates peaks required for each iteration. As the results shown in Fig. 2i, we demonstrated that the strategy can enhance the accuracy of SV peak identification, and using the HDA peaks in the initial phase can accelerate convergence. Details on ablation experiments to other parameters are available in Supplementary Note 1 and Supplementary Fig. 5. Overall, the key to advantages of Descartes lies in how to construct the graph of inter-cellular correlations from both spatial and ATAC-seq perspectives, tailored to the specific characteristics of spATAC-seq data.

### SV peaks identified by Descartes align well with spatial structure of tissues

Besides the mouse brain dataset, we also collected two datasets: (i) a dataset comprising six slices of mouse embryos, served as the mouse embryo dataset; and (ii) a dataset consisting of five slices from a mix of human and mouse tissues, served as the mixed-species dataset (Supplementary Table 1). The absence of well-defined domain labels in these datasets precludes us from utilizing the aforementioned benchmarking procedures for quantitatively assessing different methods. Here we utilized the SV peaks identified by different methods to cluster spots, and then referenced spatialPCA to evaluate the spatial clusters using three metrics: the spatial chaos score (CHAOS), the median low local inverse Simpson index (LISI), and the percentage of abnormal spots (PAS). Lower scores across these metrics signify more continuous spatial distribution of clusters, thereby indicating superior performance of the corresponding method. The detail Compared to the mouse brain dataset, both two datasets are larger-scale and impose higher demands on the robustness of methods. When identifying SV peaks, SOMDE fails to produce results within 24 hours on both datasets, while Sepal encountered errors on the mixed-species dataset. As the results showed in Fig. 3a, 3c and Supplementary Fig. 6, lower metrics indicate the clustered domains using SV peaks of Descartes are more spatially continuous and smooth, thereby demonstrating the advantages of

**Fig. 3.**
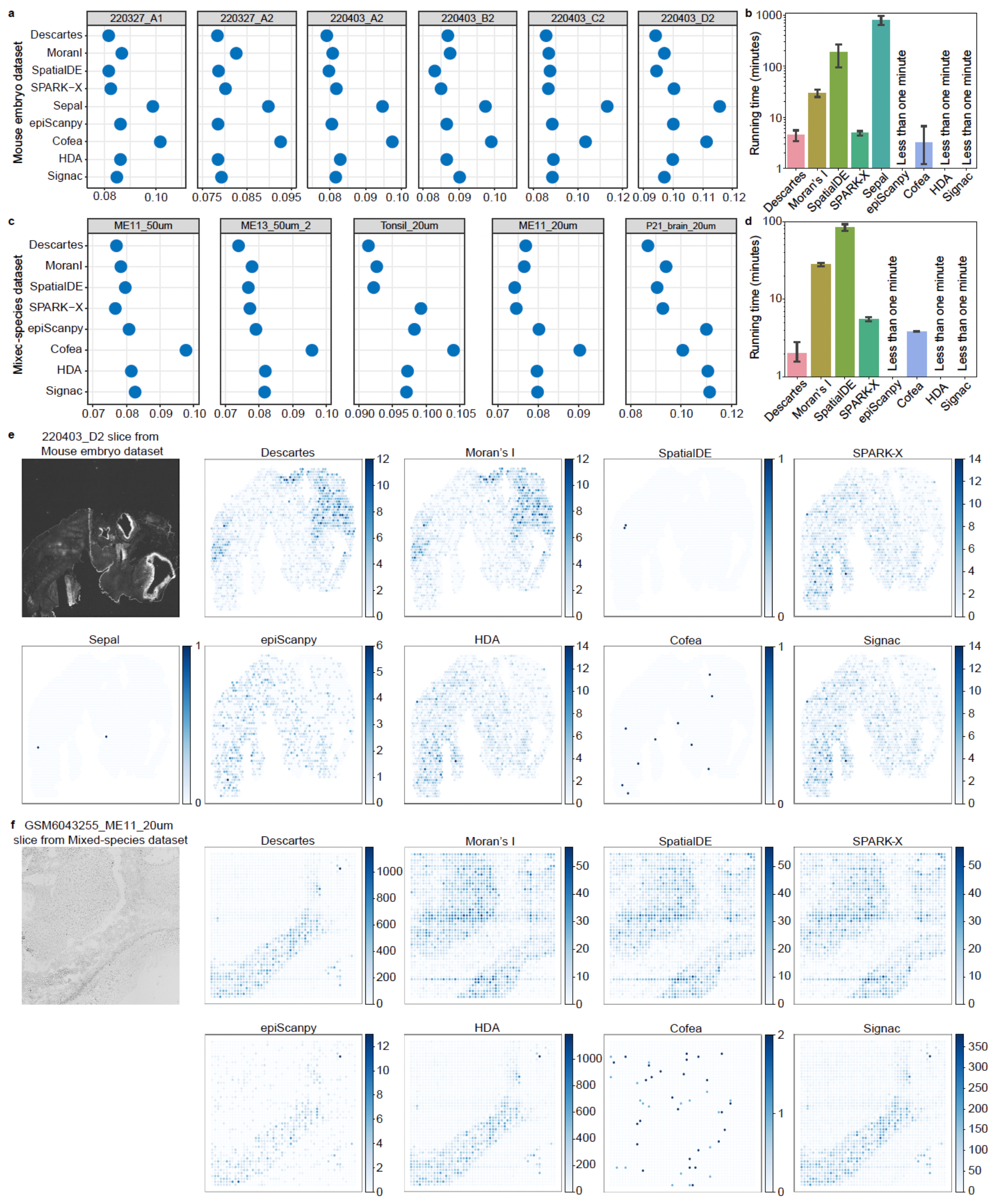
Evaluation for different methods on the mouse embryo dataset and the mixed-species dataset. **a and c**, Clustering performance evaluated by CHAOS scores using SV peaks identified by different methods on the mouse embryo dataset (**a**) and the mixed-species dataset (**c**), respectively. Due to the lack of well-annotated labels in the two datasets, we are unable to utilize label-dependent metrics for evaluation, such as NMI, ARI and AMI scores. **b and d**, Running time of different methods on the mouse embryo dataset (**b**) and the mixed-species dataset (**d**), respectively. **e-f**, The histological image (the first plot) and top-ranked SV peak identified by each method on the 220403_D2 slice from the mouse embryo dataset and the GSM6043255_ME11_20um slice from the mixed-species dataset, with the raw count values visualized in the tissue space.

Descartes in detecting spatial structure of tissues. Taking the CHAOS metric as an example, Descartes achieves the best results in 5 slices of the mouse embryo dataset and in 3 slices of the mixed-species dataset. SpatialDE gets the second-best performance in both two datasets, but the running time requires at least 10 hours and is hundreds of times longer than Descartes, underscoring a significant efficiency gap (Fig. 2b and 2d). In these datasets, methods with spatial assumptions outperform those that do not consider spatial information, consistent with observations drawn from the mouse brain dataset. We then visualized the top-ranked peaks identified by different methods and compared them with histological images (Fig. 3e and Supplementary Fig. 7 and 8). The top-ranked peaks identified by methods with spatial assumptions are generally associated with specific regions within the tissue, corroborating the insights gained from the metrics. Notably, Descartes, additionally incorporating chromatin accessibility information, identified top-ranked peaks that align well with spatial structure of tissues in all slices.

### Descartes captures cellular heterogeneity of metastatic melanoma

Next, we turned our attention to the performance of various methods on spATAC-seq data with single-cell resolution. We collected a metastatic melanoma dataset, where Russell et al. integrated their developed Slide-tags technique with scATAC-seq, facilitating simultaneous acquisition of chromatin accessibility and spatial information in individual cells^9^ (Supplementary Table 1). In their study, Russell et al. also provided well-annotated labels of cell types, allowing us to conduct a benchmarking of Descartes and baseline methods. The benchmarking is analogous to that applied to the mouse brain dataset, with the distinction that cell type labels were employed instead of domain labels. Sepal reported errors on this dataset, while SOMDE exceeded a 24-hour computation time. Benchmarking results for other methods, as illustrated in Fig. 4a (with pre-processed results in Supplementary Fig. 9), demonstrate that Descartes excels in facilitating cell clustering and revealing cell type-specific signals, thereby affirming its superiority in identifying SV peaks. Given the dataset encompassed a total of 53,431 peaks, closely aligning with our predetermined number of SV peaks, the cluster performance across different methods is relatively consistent. Note that, with the exception of two tumor subtypes, the spatial distribution patterns of various cell types are not significantly distinct (Fig. 4d), and thus methods with spatial assumptions do not show the pronounced advantages as in the mouse brain dataset. Within this dataset, Descartes also demonstrates its notable efficiency on running time over other methods with spatial assumptions (Fig. 4b). In terms of cell type-specific signals, methods that show superior performance, such as Descartes, Moran’s I, SPARK-X, epiScanpy, HDA, and Signac, primarily focus on cell types with a higher cell count (cell types are ordered by cell count from top to bottom in Fig. 4c-d). Notably, Descartes can focus on a broader range of cell types, contributing to its distinguished clustering performance. We further visualized cells, on the tissue space using actual spatial locations and the latent space using UMAP on the scATAC-seq data matrix, respectively (Fig. 4e-f). The two spaces correspond to the spatial and chromatin accessibility information of cells, serving as the foundational elements for constructing the graph in Descartes. The majority of methods do not show discernible patterns among top-ranked peaks in the two spaces. In contrast, top-ranked peak identified by Descartes is associated with two tumour subtypes, and SPARK-X with one, marking them as relatively superior methods.

**Fig. 4.**
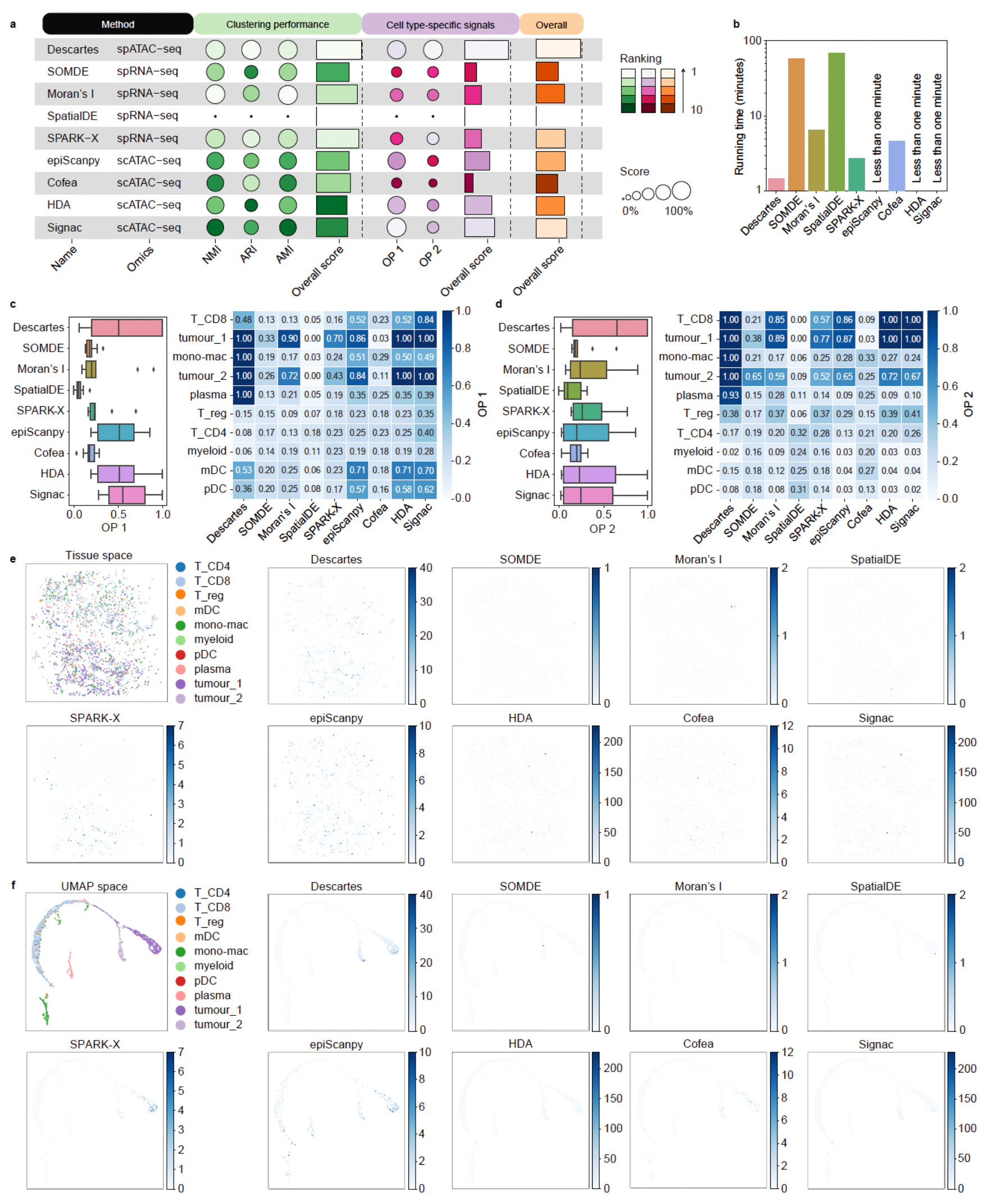
Benchmarking performance of SV peaks identification on the metastatic melanoma dataset. **a**, Overview for benchmarking results of different methods from three perspectives, that is the ability to facilitate clustering performance, capture cell type-specific signals and maintain spatial continuity (see Methods for further visualization details). **b**, Running time of different methods. **c-d**, Overlapped proportion of SV peaks identified by Descartes and baseline methods with domain-specific peaks identified by the ‘tl.rank_features’ function in epiScanpy (**c**) and “FindAllMarkers” function in Signac (**d**). **e**, The top-ranked SV peak identified by each method in the tissue space, compared to histological image (the first subplot). **f**, Top-ranked SV peak identified by each method, compared with cell type labels (the first subplot), in the UMAP space.

### Descartes imputes data using the graph of inter-cellular correlations

SpATAC-seq data typically suffer from noise and a large number of missing values, leading to the inaccuracy of downstream analysis. A key feature of Descartes is its ability to denoise data and restore missing values, by utilizing the graph-based neighbor relationships between spots. The data imputation procedure can be categorized into four cases (details in the Methods Section): (i) case 1: based on the graph of spatial locations; (ii) case 2: based on the graph of chromatin accessibility; (iii) case 3: based on the graph of inter-cellular correlations, that is the integration of case 1 and case 2; (iv) case 4: augmenting case 3 with raw data. Taking the E13_5-S1 slice of the mouse brain dataset as an example, we applied Descartes on to select 10,000 SV peaks and then performed cell clustering and uniform manifold approximation and projection (UMAP) visualization on the data before and after imputation. As shown in Fig. 5a and 5b, except for case 2, all other cases improve the accuracy of clustering, and different domains are better separated in the low-dimensional space after imputation. Using only spatial information, except for marginally enhanced spatial continuity (CHAOS), case 2 does not surpass the results using the original data in other metrics. Using only chromatin accessibility information (case 1) significantly improves clustering results, but slightly disrupts spatial continuity (LISI). In contrast, the fusion of two types of information (case 3 and 4) leads to better performance than using either type of information alone, suggesting that indispensability of both spatial and chromatin accessibility information when constructing the graph of inter-cellular correlations.

**Fig. 5.**
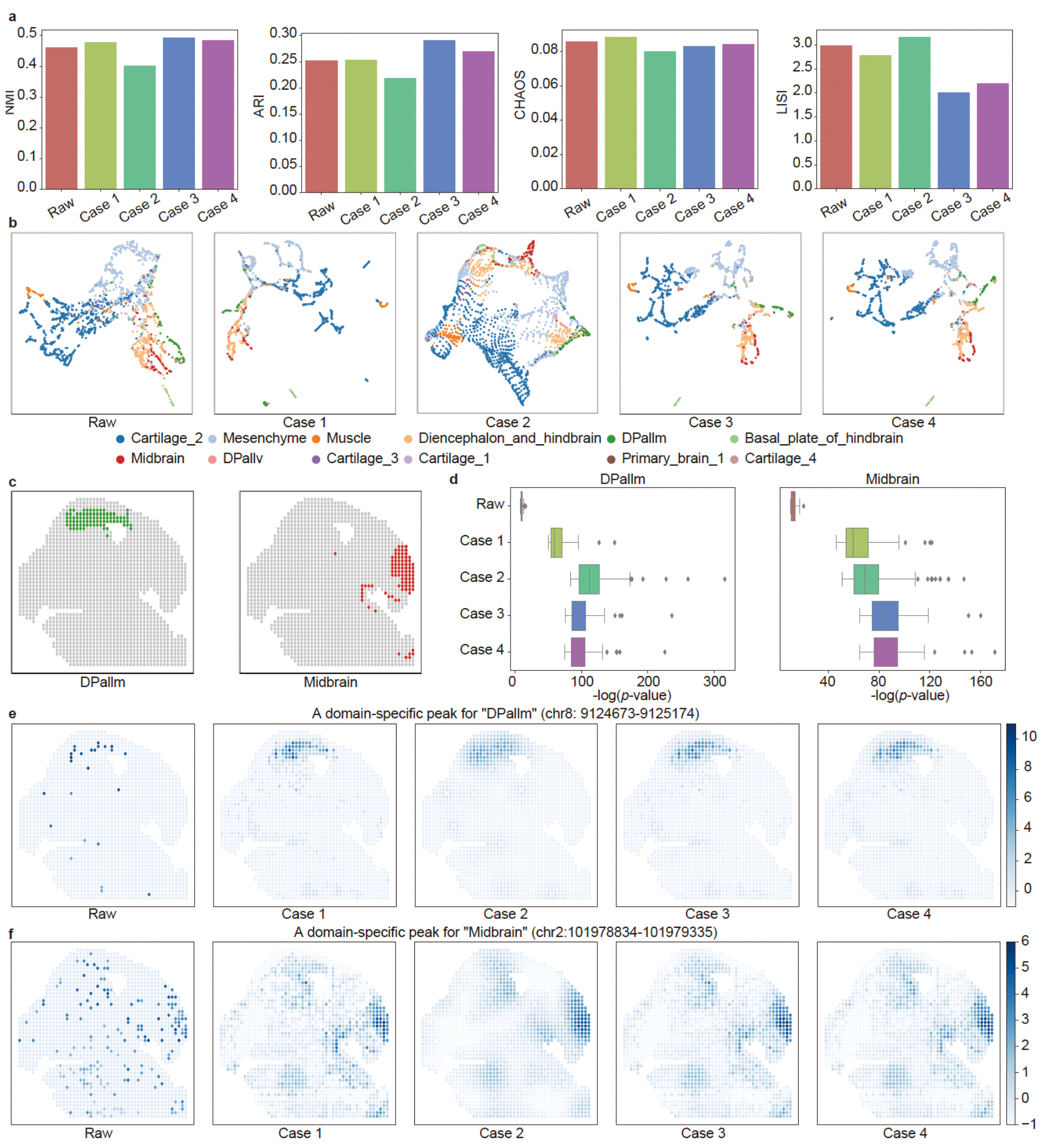
Descartes enables data imputation using the graph of inter-cellular correlations. **a**, Evaluation of clustering performance on the E13_5-S1 slice from the mouse brain dataset, assessed using NMI, ARI, CHAOS, and LISI scores. **b**, Visualization of spots in the UMAP space. **c**, Spatial locations of the domains “DPallm” and “Midbrain” in the tissue space. **d**, Statistical significance of domain-specific peaks for “DPallm” and “Midbrain”, evaluated through p-values generated by the “tl.rank_features” function in epiScanpy. **e-f**, Visualization of domain-specific peaks for “DPallm” (chr8: 9124673-9125174) (**e**) and “Midbrain” (chr8: 9124673-9125174) (**f**), in tissue space. In **a, b, d, e**, and **f**, the comparison between raw data and data imputed by Descartes is showcased. Case 1 to 4 denote as different imputation strategies implemented in Descartes (details in Methods): (i) case 1: based on the graph of spatial locations; (ii) case 2: based on the graph of chromatin accessibility; (iii) case 3: based on the graph of inter-cellular correlations, that is the integration of case 1 and case 2; (iv) case 4: augmenting case 3 with raw data.

Descartes also enables imputation in enhancing the distinct signals of different domains. Taking ‘DPallm’ and ‘Midbrain’ domains (the spatial locations are illustrated in Fig. 5c) as examples, we found that p-values (outputted by the ‘tl.rank_features’ function in epiScanpy) of domain-specific peaks significantly decreases after imputation (Fig. 5d), demonstrating the ability of Descartes in recovering signal disparities across different domains. To look deep in the differences of data imputation cases, we selected and visualized a domain-specific peak from each domain, respectively (Fig. 5e and f). Using only chromatin accessibility information (case 1) precisely boosts the signal within the domain, but disrupts spatial continuity. Using only spatial information (case 2) is akin to applying a low-pass filter to the signal in space, which enhances signals within the domain but also inadvertently amplifies out-of-domain noises. Due to the incorporation of more diverse information, Case 3 and 4 show more precise imputation effects, proving to be more applicable in real-world scenarios.

### Descartes enables peak module identification

Features with co-variation can be clustered into a module, facilitating the identification of genes or peaks specifically associated with the development of a cell type even a specific tissue structure. Utilizing modules of features for analysis, as opposed to focusing on individual features, can mitigate noise and be more efficient for researchers, making the precise identification of modules with specific patterns crucial. However, compared to gene module identification via spRNA-seq data, the field for identifying peak modules based on spATAC-seq data still remains a significant gap. Moreover, due to the high-dimensional nature of spATAC-seq data and the highly discrete patterns of chromatin accessibility in space, precise peak module identification is a formidable challenge. To fill this gap, Descartes leverages constructed graphs to calculate the similarity between peaks in terms of spatial locations and chromatin accessibility, facilitating the identification of peak modules through hierarchical clustering. In testing Descartes on the E13.5-S1 slice of the mouse brain dataset, we initially identified 10,000 SV peaks and clustered them into eight peak modules of varying sizes (Fig. 6a). Next, we averaged read counts of peaks within these modules, visualized peak modules on the tissue space, and compared them with the spatial distribution of domains and tissue structures (Fig. 6b and 6c). The results reveal that nearly every peak module corresponds to one or more specific domains: modules 1 and 3 align with the “Mesenchyme” and “Muscle” domains, respectively; module 2 correlates with peripheral regions across multiple tissues encompassing various domains, including “Basal_plate_of_hindbrain”, “Dpallm”, “Dpallv”, “Midbrain”, and “Diencephalon_and_hindbrain”; modules 4, 6, and 7 correspond to two distinct regions within the “Cartilage_2” domain; modules 5 and 8 exhibit highly similar accessible patterns, covering regions within both the “Cartilage_2” and “Muscle” domains. We then performed GREAT analysis^33^ to identify significant pathways associated with peaks from each module. As illustrated in Supplementary Fig. 10, the top-5 most significant pathways for each module are respectively associated with the development of various tissues in the mouse brain, aligning with the fetal stage of the mice used in the dataset. Furthermore, we assessed the overlap between peaks in each module and domain-specific peaks across different domains and found a strong alignment with the spatial visualization, demonstrating the potential of Descartes in peak module identification (Fig. 6d).

**Fig. 6.**
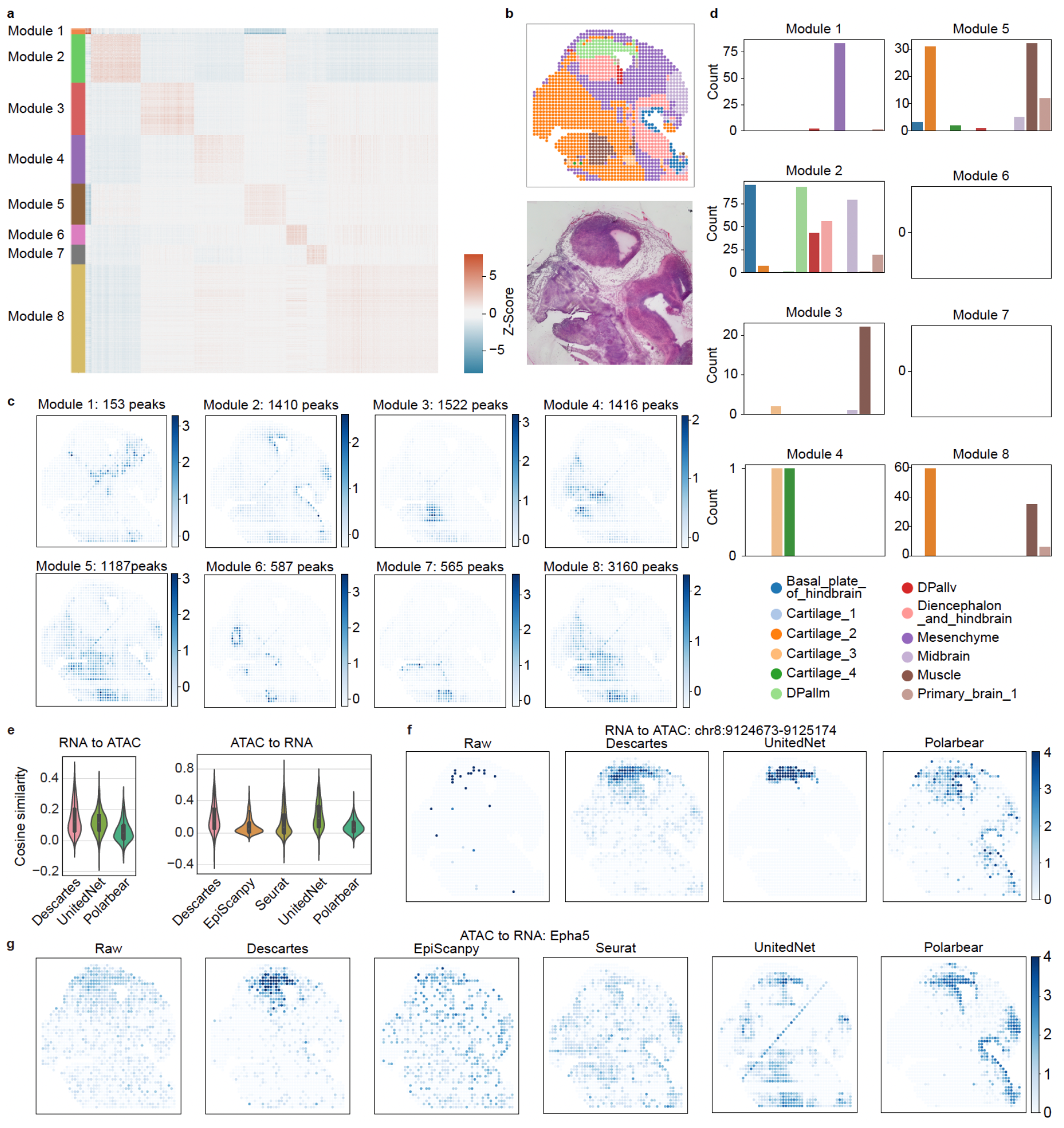
Descartes facilitates peak module identification and detection of gene-peak interaction. **a**, Heatmap of peak-peak correlations generated by Descartes. 10,000 SV peaks identified by Descartes are grouped into 8 modules. **b**, Visualization of domains in the tissue space (top) and the corresponding histological image (right). **c**, Visualization of signals for each peak module in tissue space. Signals are averaged using raw counts of peaks from each module. **d**, Overlapped counts between domain-specific peaks and peaks in each module. **e**, Cosine similarities between the raw data and predicted data by different methods in the RNA to ATAC (left) and ATAC to RNA (right) transformation task. **f-g**, Comparison between the raw data and predicted data by different methods, on the domain-specific peak (chr8: 9124673-9125174) and gene (Epha5) for “DPallm”, in the RNA to ATAC (**f**) and ATAC to RNA (**g**) translation task. The raw data shown in **f** and **g** is performed TF-IDF and z-score transformation, and the predicted data is performed z-score transformation. All experiments and subplots corresponding to the whole figure are performed on the E13.5-S1 slice of the mouse brain dataset.

### Descartes links gene expression and chromatin accessibility from spatial multi-omics data

Linking gene expression patterns to chromatin accessibility is a crucial step for constructing gene regulatory networks. leveraging the graph of inter-cellular correlations, Descartes can obtain a gene-peak correlation matrix, which captures the intensity of interactions between genes and peaks (Methods). Utilizing the E13_5-S1 slice from the mouse brain dataset as a case study, we applied the Descartes framework to identify 2,000 SV genes and 20,000 SV peaks. To assess the accuracy of gene-peak correlations identified by Descartes, we conducted a novel evaluation that uses the correlation matrix for reciprocal prediction of gene expression and chromatin accessibility and measures the performance by cosine similarity between predicted and original values. We only used the top twenty and bottom five peaks (or genes) most correlated with each gene (or peak) for prediction, with the corresponding correlation values for computation. For benchmarking, we compared two data-driven cross-omics translation methods, UnitedNet and Polarbear, as baselines, and two knowledge-based methods, functions for calculating gene activity scores in epiScanpy and Seurat. UnitedNet and Polarbear were trained and predicted on the same dataset, and such tasks are simpler than their real-world applications. As shown in Fig. 6e, except in the task transforming ATAC-seq to RNA-seq where UnitedNet excels, Descartes outperforms methods specifically designed for cross-omics prediction, suggesting the potential for elucidating gene regulatory networks from spatial multi-omics data. Knowledge-based methods (epiScanpy and Seurat), focusing only on distances between peaks and genes without updating information from training data, generally underperforms. The low cosine similarity values across methods may be attributed to differences in the sparsity of real and predicted data. Raw spatial multi-omics data, such as the domain-specific gene Epha5 and peak chr8:9124673-9125174, initially disperses in space, while the predicted data are imputed and exhibits greater spatial continuity (Fig. 6f and 6g). Visualization results demonstrate that Descartes effectively restored and enhanced the raw signals. UnitedNet, a more complex neural network-based method, precisely enhances domain-specific signals in the RNA-seq to ATAC-seq transformation task but did not perform as well in other scenarios, but, like other baseline methods, falls short in another task.

## Discussion

Recent involutions in spatial sequencing technologies can simultaneously capture spatial location and chromatin accessibility of cells, and also increase the demand for SV peaks identification tailored for modeling spATAC-seq data. In this article, we introduce Descartes, a method based on the graph of inter-cellular correlations, for identifying peaks characterized by both spatial variation and cellular heterogeneity. To our best knowledge, Descartes is the first method for SV peaks identification tailored for spATAC-seq data. To deal with the challenge posed by the overly discrete spatial patterns of chromatin accessibility in spATAC-seq data, Descartes employs following strategies to capture precise neighborhood relationships among cells: (i) integrating the graph constructed from chromatin accessibility information into the spatial graph; (ii) considering a broad range of neighboring cells when constructing the spatial graph, and (iii) iteratively updating the graph built from chromatin accessibility information with each iteration of SV peaks. To comprehensively evaluate our method, our benchmarking pipeline spans 16 slices from 4 datasets and incorporates three aspects from enhancing clustering performance, capturing domain-specific signals, and preserving spatial continuity. The benchmarking results demonstrate that Descartes surpasses other methods with only spatial assumptions in both the accuracy and efficiency of SV peak identification. Due to the lack of spatial information, methods based on scATAC-seq data failed to maintain spatial continuity in the identified SV peaks, thereby undermining the accuracy of domain identification. Through case studies on the E13.5-S1 slice of the mouse brain dataset, we demonstrate the potential of Descartes for data imputation, peak module identification and gene-peak interaction detection. Overall, Descartes offers an effective and valuable tool for spATAC-seq data analysis, contributing to detection of spatial chromatin accessibility patterns with inter-cellular correlations.

Despite the progress achieved so far, Descartes still has several potential directions that need to be improved. First, to ensure high-efficiency of Descartes on large-scale datasets, we will introduce downsampling or self-organizing maps for keeping the number of nodes in the constructed graph within a manageable range. The key concept of Descartes is based on the graph of inter-cellular correlations, and its computational complexity scales quadratically with the number of cells. As spATAC-seq technologies evolve, increasing the spots (or cells) within datasets, Descartes may face efficiency challenges. Second, we aim to refine existing simulation methods, such as simCAS, to provide multi-scenario simulated spATAC-seq data for systematic evaluation. Most feature selection methods based on spRNA-seq and scATAC-seq leverage simulated data for evaluation, as such data typically come with precise annotations. Finally, considering the availability of other spatial omics data, such as spatial metabolic data^34^ and spatial CITE-seq data^35^, we will broaden the application scope of Descartes in our future work.

## Methods

### Construction for graph of spatial locations

To capture spatial accessibility pattern of each peak, Descartes construct a spatial graph based on spatial locations of spots (can also be replaced by cells). We suppose that ***X*** = (***x***_.*1*_, *…*, ***x***_.*N*_) denotes as spatial locations of all *N* spots in a single slice, and the coordinates of the spatial locations are typically two-dimensional (i.e., ***x***_.*i*_ can be represented as (*x*_*i1*_, *x*_*i2*_)^*T*^). Descartes first calculates the mean Euclidean distance between each spot and its nearest neighbor, denoting it as the standard distance 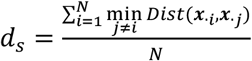, and then connects each spot to its neighbors within five times *d*_*s*_ to obtain a spatial graph. The edge weight of the spatial graph is inversely proportional to the Euclidean distance between spots, i.e., the edge weight between spot *i* and spot *j* is given by

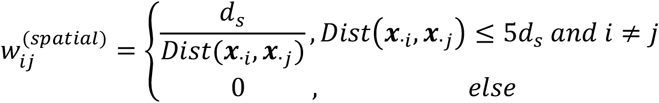

where 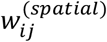 is an element in edge weight set ***W***^*(spatial)*^ of the spatial graph, and *Dist*(***x***_.*i*_, ***x***_.*j*_) denotes the Euclidean distance between spot *i* and spot *j* in tissue space.

### Construction for graph of chromatin accessibility

The majority of graph-based existing methods for selecting primarily rely on spatial positional information for graph construction, while the constructed graph tends to overlook the inherent spot-spot relationships associated with gene expression or chromatin accessibility. In contrast to those methods, Descartes incorporates a graph constructed from the ATAC-seq matrix and integrates it with the spatial graph. Due to the characteristics of high dimensionality, sparsity, and noise in the ATAC-seq matrix, constructing a graph directly from the raw count matrix is deemed impractical. Assuming that ***Y*** ∈ ℝ^*MxN*^ denotes as the raw peak-by-spot (or peak-by-cell) count matrix with *N* spots and *M* peaks, Descartes first applies term frequency-inverse document frequency (TF-IDF) transformation as in Signac^29^ to obtain a continuously valued matrix (denoted as 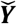). Then Descartes selects a subset of peaks (50,000 peaks as the default) and performs PCA transformation to obtain a PC-by-spot matrix ***P*** = (***p***_.*1*_, *…*, ***p***_.*N*_) ∈ ℝ^*10×N*^. Note that as an iterative method, in the initial iteration, Descartes directly selects peaks based on their decreasing order of accessible degree, and in subsequent iterations, involves the selection according to the ranking from the previous iteration. With the PC-by-spot matrix ***P***, Descartes connects each spot to its 20 nearest neighbors and defines the edge weight between spot *i* and spot *j* as

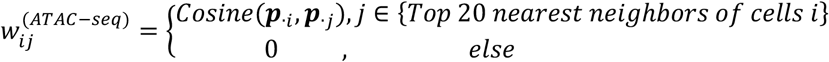

where *Cosine(****p***_*i*_, ***p***_*j*_) denotes as the cosine similarity between vector ***p***_*i*_ and ***p***_*j*_, 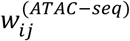 is an element in edge weight set ***W***^*(ATAC−seq)*^ of the chromatin accessibility graph. **SV peaks selection**. Given the graph of spatial locations and the graph of chromatin accessibility, Descartes directly integrates the edges of them to obtain a graph of inter-cellular correlations. The edge weight of spot *i* and spot *j* in the newly-formed graph can be calculated as

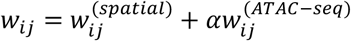

where *α* is the factor for balancing the two types of edge weights (1.5 in default), and *w*_*ij*_ is an element in edge weight set ***W*** of the inter-cellular correlations graph. On the other hand, Descartes perform z-score transformation on the matrix 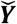 to obtain the matrix 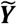. The key concept of Descartes is grounded in that peaks with regional continuity in the graph will be identified as informative peaks (i.e., SV peaks), while peaks with disparate accessibility levels between the two spots connected by the majority of graph edges will be considered as non-informative peaks (i.e., peaks that need to be filtered out). Under the concept, Descartes evaluated each peak by an importance score using self-correlations, which can be obtained by

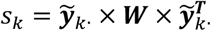

where 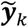. is a row vector in the matrix 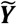 representing the peak *k*, and *s*_*k*_ is the importance score of the peak *k*. Based on the importance score, Descartes sorts all peaks and feeds the ranking back into the step of constructing the graph of chromatin accessibility. The iterative process continues until the ranking stabilizes. Descartes involves four iterations as the default setting, with outputting the final iteration’s importance scores as the ultimate results. It is noteworthy that after performing omics-specific preprocessing and transforming the values of each feature into z-score-transformed values, Descartes can also be employed to obtain the importance scores of features for any type of spatial sequencing data. Researchers can utilize the importance scores to fit various distributions to obtain p-values for all peaks. Alternatively, they can straightforwardly designate a specific number or a predetermined proportion of peaks as SV peaks.

### Peak module identification

Based on the graph of inter-cellular correlations and its edge weights, Descartes can obtain the peak-peak similarity matrix ***S***^*(peak-peak)*^ by

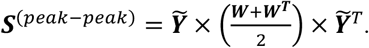

Subsequently, Descartes transforms the matrix ***S***^*(peak-peak)*^ to a peak-peak distance matrix ***D***^*(peak-peak)*^ through (i) being subtracted by the 99.5th percentile of the elements in ***S***^*(peak-peak)*^ from each element in ***S***^*(peak-peak)*^ and (ii) setting the diagonal elements and negative values of the resulting matrix to zero. Finally, Descartes utilizes the scipy package for hierarchical clustering (with the method parameter set to ‘ward’) to identify peak modules.

### Gene-peak interaction detection

For transcriptomics data within spatial multi-omics data, Descartes uses the same preprocessing procedure as in Seurat and Scanpy. Specifically, Descartes (i) scales the library size of each spot to 10,000, (ii) performs log*(x* + *1)* transformation and (iii) performs z-score transformation, on the raw gene-by-cell (or gene-by-spot) matrix. Then Descartes integrates the processed matrix with the z-score-transformed peak-by-spot matrix to obtain a feature-by-spot matrix 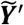, where features encompass all genes and peaks. Using the same calculation formula as for ***S***^*(peak-peak)*^, Descartes can derive a correlation matrix among all features, from which the corresponding submatrix reveals the strength of gene-peak or peak-gene interactions. **Data imputation**. By incorporating the chromatin accessibility information of each spot with that of other cells, Descartes enables data imputation on the raw data. Specifically, utilizing the graph of inter-cellular correlations and the weights of its edges, Descartes performs a weighted averaging of the data from nearest neighbor spots, and adds the result onto the raw data, that is

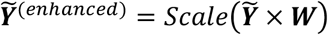

where *Scale(*.) denotes the function of z-score transformation, and 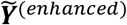 represents the enhanced matrix that can be directly used for downstream analysis. Descartes also provides two alternatives: (i) imputing data only based on neighborhood relationships from the spatial graph or the chromatin accessibility graph, that is substituting ***W*** in the formula for obtaining 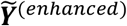 with ***W***^*(spatial)*^ or ***W***^*(CAS)*^, and (ii) incorporating the original data with the imputed data, that is 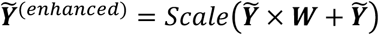

### Data collection

The mouse brain dataset^8^, which simultaneously provides capture chromatin accessibility and gene expression of spots, consists of four tissue slices from mouse fetal brain at stage E11.0, E13.5, E15.5 and E18.5, respectively. In addition, the dataset also provides accurate manual anatomical annotations which reveal major tissue organizations of these slices, with reference to Kaufman’s Atlas of Mouse Development and Allen Brain Atlas. The mouse embryo^6^ dataset composed of six tissue sections from three stages of mouse gestational development, including E12.5, E13.5 and E15.5 embryonic days. The mixed-species dataset^7^ comprises a collection of tissues from human and mouse, including five slices, including five slices—three from mouse embryos, one from the mouse brain, and another from human tonsil. The metastatic melanoma dataset is a single-cell resolution dataset, also serving as a spatial multi-omics dataset^9^. In this dataset, Russell et al. have ingeniously integrated their Slide-tags technology with scRNA-seq and scATAC-seq, allowing for the simultaneous capture of chromatin accessibility and gene expression at the cellular level. Besides, Russell et al. also provided well-annotated cell type labels of the dataset in their study. A summary of the above datasets is shown in Supplementary Table 1.

### Model evaluation

#### SV peaks selection

Due to the limited knowledge about the congruent relationship between domains (or cell types) and peaks, systematically benchmarking methods for SV peaks selection remains a tough challenge. For datasets with well-annotated labels, such as the mouse brain and metastatic melanoma datasets, we follow and improve the quantitative evaluation procedure from our prior research^27^, that is evaluating whether SV peaks identified by Descartes can (i) perform better in facilitating cell clustering and (ii) capture more domain-specific (or cell type-specific) signals. The design of the evaluation framework and metric visualization is also informed by scIB^32^, with specific calculations detailed as follows.

From the perspective of facilitating cell clustering, we utilize different methods to select SV peaks, and then perform unsupervised clustering to obtain annotated spatial domains. Due to the absence of analytic methods tailored for spATAC-seq data, we resort to the Signac^29^ to obtain low-dimensional representations and clustering labels of spots (or cells). When clustering spots by the leiden clustering (the default clustering method in Signac), we use a binary search to ensure that the number of clusters matches the number of domain labels. The clustering accuracy is assessed by NMI, ARI, and AMI scores, and higher values of the three metrics indicate that the method retains more spatial domain information in the identified SV peaks.

From the perspective of capturing domain-specific signals, we assess the performance using the overlap proportion between identified SV peaks and domain-specific accessible peaks. Specifically, given the domain labels in a specific dataset, we first use “FindAllMarkers” function in Signac^29^ or the ‘tl.rank_features’ function in epiScanpy^28^ to extract 100 domain-specific peaks for each domain, and then compare the OP between specific peaks for each domain and SV peaks identified by various methods. OP1 corresponds to the overlap using the “FindAllMarkers” function in Signac, and OP2 corresponds to the overlap using the ‘tl.rank_features’ function in epiScanpy. A higher OP value indicates that SV peaks encompass more heterogeneity information related to specific domains.

Referencing scIB, to ensure that each metric contributes equally to each perspective and possesses the same discriminative power, we perform min-max scaling on individual metrics among different methods. Specifically, for each metric, we scale the highest value among different methods into 1 and the lowest value into 0. After scaling, all metrics of each method for each perspective is averaged to obtain a score, with a higher value indicating superior performance of the method in that perspective. The overall score of each method is then determined by averaging the scores of the two perspectives, serving as the final measure of each method’s performance.

For the mouse embryo and mixed species datasets, the absence of well-annotated labels precludes the same quantitative assessment as aforementioned. We here assess SV peaks identified by each method based on the ability to preserve spatial continuity as in spatialPCA^36^. Specifically, we perform cell clustering based on SV peaks, and then use CHAOS, median LISI, and PAS scores to measure the spatial continuity of the clusters. Lower values of the three metrics indicate better performance of the corresponding method. The workflow for cell clustering is consistent with the aforementioned setting, and the number of clusters is set to 10 for all slices.

Detailed formulas for all the metrics mentioned above are provided in Supplementary Note 2. In addition to assessing the accuracy of SV peak identification, we conducted evaluations of the running time for different methods, i.e., the time they require from initial data processing to obtaining the ranking of each peak. All experiments were conducted on a server with 128GB of memory and equipped with 32 units of 13th Gen Intel(R) Core(TM) i9-13900K.

#### Baseline methods

Given the absence of methods specifically designed for spATAC-seq data, we assessed the performance of Descartes against two categories of methods: (i) those based on spRNA-seq data, including SOMDE^15^, SpatialDE2^11^, SpatialDE^12^, SPARK-X^13^, SPARK^31^, scGCO^14^, Sepal^16^, and Moran’s I^17^; and (ii) those based on scATAC-seq data, including Cofea^27^, HDA^22-26^, Signac^29^, and epiScanpy^28^. HDA, notably the most commonly used feature selection method for scATAC-seq data analysis, selects peaks with at least one read count in a majority of cells. We implemented HDA in Python following its instruction. EpiScanpy and Signac, both important pipelines for scATAC-seq data processing, were utilized solely for the feature selection functions as baseline comparison methods. The implementation of other methods was conducted using source code and default parameters provided in their respective studies.

## Data availability

The mouse brain dataset^8^ is available accessed under National Genomics Data Center accession number (OEP003285, www.biosino.org/node/project/detail/OEP003285). The mouse embryo dataset^6^ can be accessed from GEO under accession number GSE214991. The mixed-species dataset^7^ is deposited in the Gene Expression Omnibus (GEO) with accession code GSE171943. The metastatic melanoma dataset^9^ is available at the Gene Expression Omnibus under accession number GSE244355.

## Code availability

The Descartes software, including detailed documents and tutorial, is freely available on GitHub under a MIT license (https://github.com/likeyi19/Descartes), with the version used in the manuscript also deposited in Zenodo (https://zenodo.org/records/10774503).

## References

1. Zhang, M. et al. Spatially resolved cell atlas of the mouse primary motor cortex by MERFISH. Nature 598, 137–143 (2021).

2. Asp, M. et al. A Spatiotemporal Organ-Wide Gene Expression and Cell Atlas of the Developing Human Heart. Cell 179, 1647–1660 e1619 (2019).

3. Wang, X. et al. Three-dimensional intact-tissue sequencing of single-cell transcriptional states. Science 361 (2018).

4. Rodriques, S.G. et al. Slide-seq: A scalable technology for measuring genome-wide expression at high spatial resolution. Science 363, 1463–1467 (2019).

5. Lomakin, A. et al. Spatial genomics maps the structure, nature and evolution of cancer clones. Nature 611, 594–602 (2022).

6. Llorens-Bobadilla, E. et al. Solid-phase capture and profiling of open chromatin by spatial ATAC. Nat Biotechnol 41, 1085–1088 (2023).

7. Deng, Y. et al. Spatial profiling of chromatin accessibility in mouse and human tissues. Nature 609, 375–383 (2022).

8. Jiang, F. et al. Simultaneous profiling of spatial gene expression and chromatin accessibility during mouse brain development. Nat Methods 20, 1048–1057 (2023).

9. Russell, A.J.C. et al. Slide-tags enables single-nucleus barcoding for multimodal spatial genomics. Nature 625, 101–109 (2024).

10. Palla, G., Fischer, D.S., Regev, A. & Theis, F.J. Spatial components of molecular tissue biology. Nat Biotechnol 40, 308–318 (2022).

11. Kats, I., Vento-Tormo, R. & Stegle, O. SpatialDE2: fast and localized variance component analysis of spatial transcriptomics. Biorxiv, 2021.2010. 2027.466045 (2021).

12. Svensson, V., Teichmann, S.A. & Stegle, O. SpatialDE: identification of spatially variable genes. Nat Methods 15, 343–346 (2018).

13. Zhu, J., Sun, S. & Zhou, X. SPARK-X: non-parametric modeling enables scalable and robust detection of spatial expression patterns for large spatial transcriptomic studies. Genome Biol 22, 184 (2021).

14. Zhang, K., Feng, W. & Wang, P. Identification of spatially variable genes with graph cuts. Nat Commun 13, 5488 (2022).

15. Hao, M., Hua, K. & Zhang, X. SOMDE: a scalable method for identifying spatially variable genes with self-organizing map. Bioinformatics 37, 4392–4398 (2021).

16. Andersson, A. & Lundeberg, J. sepal: identifying transcript profiles with spatial patterns by diffusion-based modeling. Bioinformatics 37, 2644–2650 (2021).

17. Palla, G. et al. Squidpy: a scalable framework for spatial omics analysis. Nat Methods 19, 171–178 (2022).

18. DeTomaso, D. & Yosef, N. Hotspot identifies informative gene modules across modalities of single-cell genomics. Cell Syst 12, 446–456 e449 (2021).

19. Wu, Y. et al. Highly Regional Genes: graph-based gene selection for single-cell RNA-seq data. J Genet Genomics 49, 891–899 (2022).

20. Chen, C., Kim, H.J. & Yang, P. Evaluating spatially variable gene detection methods for spatial transcriptomics data. Genome Biol 25, 18 (2024).

21. Charitakis, N. et al. Disparities in spatially variable gene calling highlight the need for benchmarking spatial transcriptomics methods. Genome Biol 24, 209 (2023).

22. Chen, X. et al. Cell type annotation of single-cell chromatin accessibility data via supervised Bayesian embedding. Nat Mach Intell 4, 116–126 (2022).

23. Chen, S. et al. RA3 is a reference-guided approach for epigenetic characterization of single cells. Nat Commun 12, 2177 (2021).

24. Zamanighomi, M. et al. Unsupervised clustering and epigenetic classification of single cells. Nat Commun 9, 2410 (2018).

25. Xiong, L. et al. SCALE method for single-cell ATAC-seq analysis via latent feature extraction. Nat Commun 10, 4576 (2019).

26. Liu, Q., Chen, S., Jiang, R. & Wong, W.H. Simultaneous deep generative modeling and clustering of single cell genomic data. Nat Mach Intell 3, 536–544 (2021).

27. Li, K. et al. Cofea: correlation-based feature selection for single-cell chromatin accessibility data. Brief Bioinform 25 (2023).

28. Danese, A. et al. EpiScanpy: integrated single-cell epigenomic analysis. Nat Commun 12, 5228 (2021).

29. Stuart, T., Srivastava, A., Madad, S., Lareau, C.A. & Satija, R. Single-cell chromatin state analysis with Signac. Nat Methods 18, 1333–1341 (2021).

30. Moran, P.A. Notes on continuous stochastic phenomena. Biometrika 37, 17–23 (1950).

31. Sun, S., Zhu, J. & Zhou, X. Statistical analysis of spatial expression patterns for spatially resolved transcriptomic studies. Nat Methods 17, 193–200 (2020).

32. Luecken, M.D. et al. Benchmarking atlas-level data integration in single-cell genomics. Nat Methods 19, 41–50 (2022).

33. McLean, C.Y. et al. GREAT improves functional interpretation of cis-regulatory regions. Nat Biotechnol 28, 495–501 (2010).

34. Sun, C. et al. Spatially resolved multi-omics highlights cell-specific metabolic remodeling and interactions in gastric cancer. Nat Commun 14, 2692 (2023).

35. Liu, Y. et al. High-plex protein and whole transcriptome co-mapping at cellular resolution with spatial CITE-seq. Nat Biotechnol 41, 1405–1409 (2023).

36. Shang, L. & Zhou, X. Spatially aware dimension reduction for spatial transcriptomics. Nat Commun 13, 7203 (2022).

