## Supplementary Notes, Figures and Tables for "Detection of spatial chromatin accessibility patterns with inter-cellular correlations"

**Supplementary Note 1: Ablation experiments of Descartes on the mouse brain dataset**

We utilized four well-annotated slices from the mouse brain dataset to examine the impact of different parameters in Descartes on the accuracy of SV peaks identification. We varied each hyperparameter in Descartes in turn and fixed the other hyperparameters to the default setting, to obtain different variants of Descartes. Using 10,000 SV peaks identified by Descartes and its variants for clustering, we quantitively assessed the impact of different parameters with NMI scores as the metric. Beyond the ablation experiments about key parameters or modules presented in the main text, here the analysis primarily focused on the following four aspects concerning their impact on Descartes: (i) the number of neighboring cells (or spots) considered when constructing the graph of chromatin accessibility; (ii) the number of principal components (PCs) retained after PCA when constructing the graph of chromatin accessibility; (iii) the strategies of calculating cell-cell similarity and edge weights when constructing the graph of chromatin accessibility; (iv) the relationship between edge weights and cell-cell distance when constructing the graph of spatial locations. The first three aspects pertain to the graph of chromatin accessibility, while the last is related to the graph of spatial locations.

Results related to (i) are shown in Supplementary Fig. 5a, indicating that the number of neighboring cells considered during the construction of the chromatin accessibility graph does not significantly impact performance, with the optimal results occurring at 20 neighboring cells. Such the result is attributed to the iterative strategy in Descartes for graph construction, which accurately captures the precise neighboring relationships of cells. The optimal number of neighbors is thus more about averaging cells of different domains.

For (ii), as illustrated in Supplementary Fig. 5b, despite the conventional belief that higher dimensions of low-dimensional representations can preserve more information on cellular heterogeneity, we found that selecting either 5 or 10 PCs yielded better results. The paradox arises because the variance captured by additional PCs decreases, thus diminishing the effectiveness in capturing cellular heterogeneity and making the similarities or distances in the low-dimensional space less reflective of the real inter-cellular correlations.

Regarding (iii), we compared four scenarios: using cosine similarity for neighboring point similarity and edge weighting (Cosine); using inverse Euclidean distance for neighbor similarity and Jaccard coefficient for edge weighting (Euclidean+Jaccard); using cosine similarity for neighbor similarity and Jaccard coefficient for edge weighting (Cosine+Jaccard); and using Pearson similarity for both neighbor similarity and edge weighting (Pearson). The comparison of these scenarios (Supplementary Fig. 5c) revealed that cosine similarity more accurately delineates inter-cellular correlations in the low-dimensional space, thereby aiding in the accuracy of SV peaks identification.

For (iv), we examined three relationships between edge weights and cell-cell distance: constant, inverse, and inverse square relationships. The results, as depicted in Supplementary Fig. 5d, demonstrate that the inverse relationship more accurately captures the spatial accessibility patterns of peaks. Comparatively, a constant value is even more effective than an inverse square relationship, suggesting that the spatial accessibility patterns are highly discrete and imposing too strict a distance constraint could overlook many SV peaks. However, necessary degree of distance constraint is still required, which explains why the inverse relationship yields the best results.

**Supplementary Note 2: Detailed formulas for all the metrics**

We denote $\boldsymbol{U}$ as the ground-truth labels, $\boldsymbol{V}$ as the predicted cluster labels, and the NMI score can be calculated by the following formula:

$$NMI=\frac{MI\left( \boldsymbol{U},\boldsymbol{V} \right)}{\sqrt{H\left( \boldsymbol{U} \right)\cdot H\left( \boldsymbol{V} \right)}}$$

where $\mathrm{MI}\left( \cdot,\cdot\right)$ is used to compute the mutual entropy, and $H\left( \cdot\right)$ is used to compute the entropy.

The AMI score can be calculated as follows:

$$AMI=\frac{MI\left( \boldsymbol{U},\boldsymbol{V} \right)-E\left( MI\left( \boldsymbol{U},\boldsymbol{V} \right) \right)}{avg\left( H\left( \boldsymbol{U} \right),H\left( \boldsymbol{V} \right) \right)-E\left( MI\left( \boldsymbol{U},\boldsymbol{V} \right) \right)}$$

where $E\left( \cdot\right)$ is the expectation function. We then suppose that $u_{i}$ is the $i$th true label, $v_{j}$ is the $j$th predicted label, $n_{ij}$ is the number of spots simultaneously belonging to $u_{i}$ and $v_{j}$, $n_{i\cdot}$ is the number of spots that belong to $u_{i}$, $n_{\cdot j}$ is the number of spots that belong to $v_{j}$, and $n$ is the total number of spots. The ARI score is calculated as follows:

$$ARI=\frac{\sum_{i,j} \left( \begin{aligned} n_{ij} \\ 2 \end{aligned} \right)-\left[ \sum_{i} \left( \begin{aligned} n_{i\cdot} \\ 2 \end{aligned} \right)\sum_{j} \left( \begin{aligned} n_{\cdot j} \\ 2 \end{aligned} \right) \right]/\left( \begin{aligned} n \\ 2 \end{aligned} \right)}{\frac{1}{2}\left[ \sum_{i} \left( \begin{aligned} n_{i\cdot} \\ 2 \end{aligned} \right)+\sum_{j} \left( \begin{aligned} n_{\cdot j} \\ 2 \end{aligned} \right) \right]-\left[ \sum_{i} \left( \begin{aligned} n_{i\cdot} \\ 2 \end{aligned} \right)\sum_{j} \left( \begin{aligned} n_{\cdot j} \\ 2 \end{aligned} \right) \right]/\left( \begin{aligned} n \\ 2 \end{aligned} \right)}$$

In the computation of OP 1 and OP 2, we utilized epiScanpy (for OP 1) and Signac (for OP 2) to identify domain-specific peaks. Subsequently, we assessed the overlapped proportion between peaks identified by various methods and the domain-specific peaks. Denoting $\boldsymbol{P}$ as set of peaks selected by different methods, and $\boldsymbol{Q}$ as the set of domain-specific peaks identified by epiScanpy or Signac. The calculation formula is as follows:

$$OP 1 \left( or OP 2 \right)=\frac{\left[ \boldsymbol{P}\cap\boldsymbol{Q} \right]}{\left[ \boldsymbol{P} \right]}$$

where $\left[ \cdot\right]$ denotes the number of elements in the set.

CHAOS calculation involves two steps. Firstly, constructing a one-nearest-neighbor (1NN) graph for the spots in each spatial cluster by connecting each spot with its nearest neighbor. The edge weight between spot $i$ and spot $j$ is calculated as:

$$w_{kij}=\left\{ \begin{aligned} d_{ij}, spot i and j are connected in the 1NN graph in cluster k \\ 0, otherwise \end{aligned} \right.$$

where $d_{ij}$ is the Euclidean distance between the two spots. Then CHAOS is calculated as the mean length of the graph edges in the 1NN graph:

$$CHAOS=\frac{\sum_{k=1}^{K} \sum_{i,j}^{n_{k}} w_{kij}}{N}$$

where $N$ denotes the number of spots; $K$ denotes the number of domains; and $n_{k}$ is the number of spots in $k$-th spatial domain. A smaller CHAOS score indicates enhanced spatial continuity.

The LISI score represents the effective number of spatial domain labels within the local spatial neighborhood of a spot, and it is calculated as:

$$S=\frac{1}{\sum_{k=1}^{K} p\left( k \right)}$$

where $p\left( k \right)$ denotes the probability of the spatial domain cluster label being present in the local neighborhood, and $K$ represents the total number of spatial domains. We employ the median LISI score across all spots as the metric. A lower LISI score suggests a more homogeneous distribution of spatial domain clusters in the neighborhood of the spot.

The PAS score evaluates the dispersion of spots beyond their clustered spatial domain. It quantifies the proportion of spots possessing a cluster label distinct from at least six out of ten neighboring spots. A low PAS score signifies homogeneity among spots within spatial clusters.

**Supplementary Figures**

**Supplementary Figure 1**

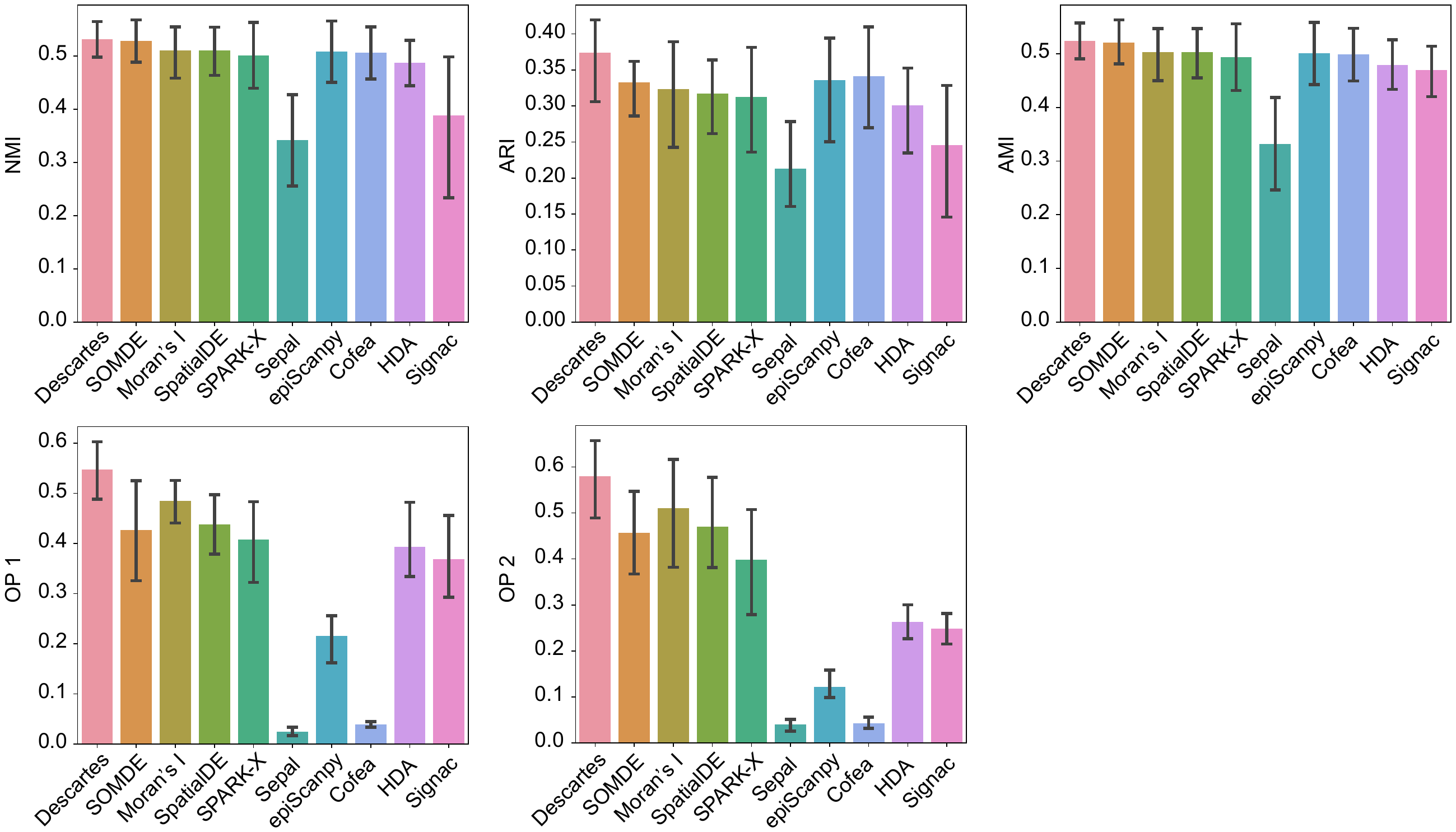

**Supplementary Figure 1.** Benchmarking results before metric transforming of different methods on the mouse brain dataset from two perspectives, that is the ability to facilitate clustering performance and capture domain-specific signals. For the perspective of facilitating clustering performance, we use NMI, ARI and AMI scores as metrics. For the perspective of capturing domain-specific signals, we use OP1 and OP2 scores as metrics. The error bars denote the 95% confidence interval, and the centres of the error bars denote the average value.

**Supplementaery Figure 2**

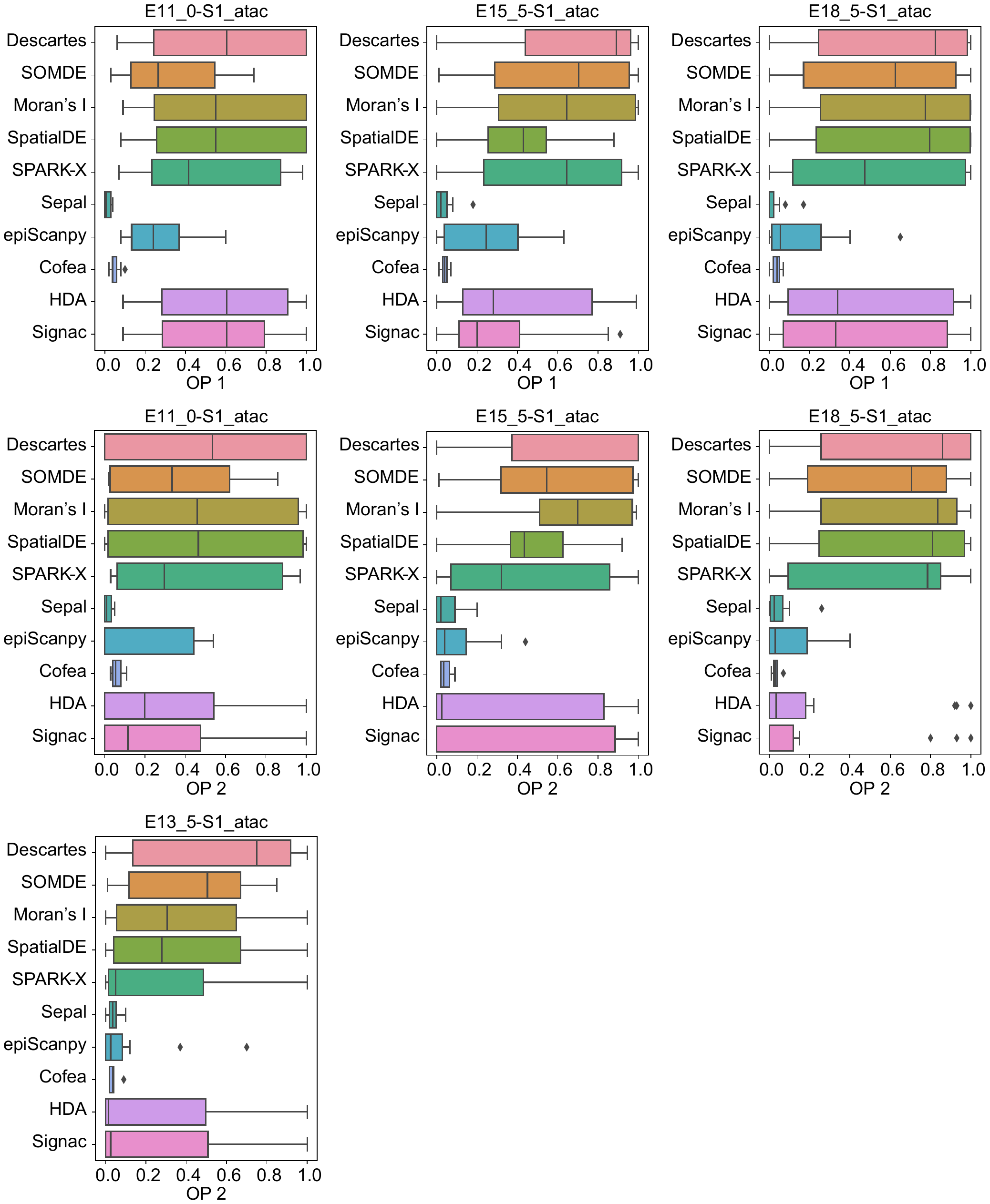

**Supplementary Figure 2.** Overlapped proportion (OP) of SV peaks identified by Descartes and baseline methods with domain-specific peaks related to overall domains. OP1 and OP2 correspond to overlaps identified by “tl.rank_features” function in epiScanpy and “FindAllMarkers” function in Signac, respectively.

**Supplementary Figure 3**

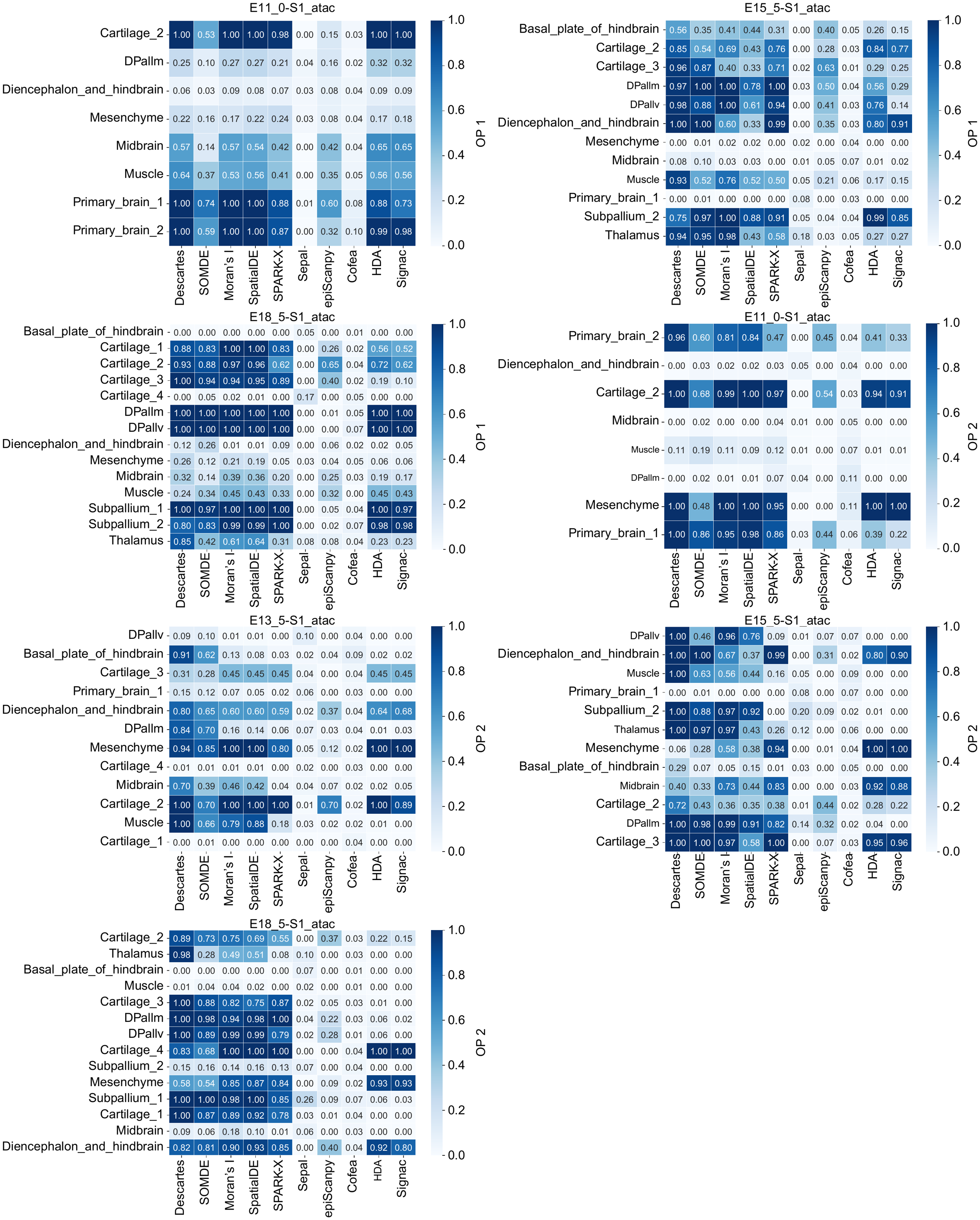

**Supplementary Figure 3.** Overlapped proportion (OP) of SV peaks identified by Descartes and baseline methods with domain-specific peaks related to each domain. OP1 and OP2 correspond to overlaps identified by “tl.rank_features” function in epiScanpy and “FindAllMarkers” function in Signac, respectively.

**Supplementary Figure 4**

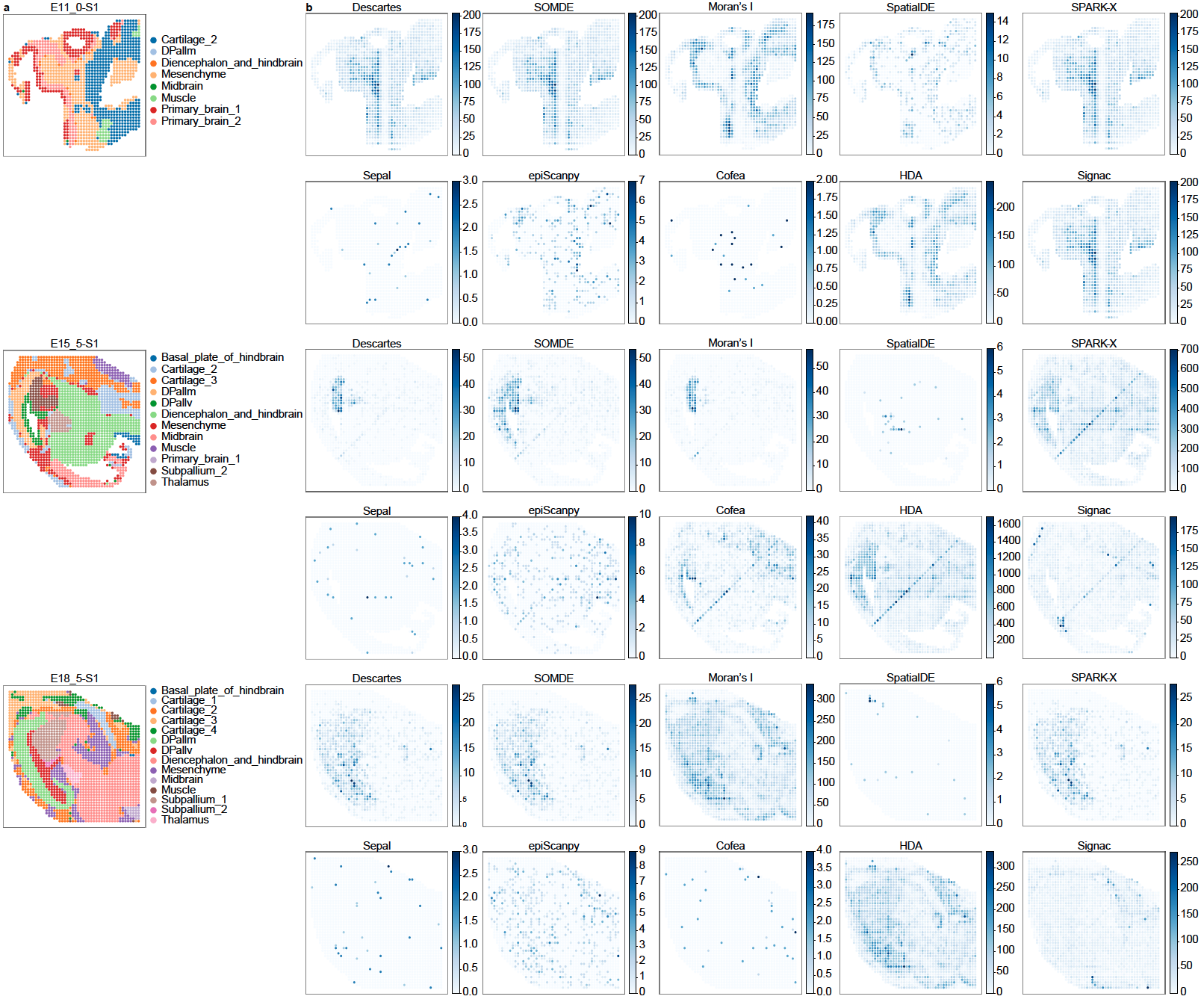

**Supplementary Figure 4. a**, Visualization of domains within the tissue space (top) and the corresponding histological images (bottom) on the slices from the mouse brain dataset. **b**, Top-ranked SV peak identified by each method on the slices from the mouse brain dataset, with the raw count values visualized in the tissue space.

**Supplementary Figure 5**

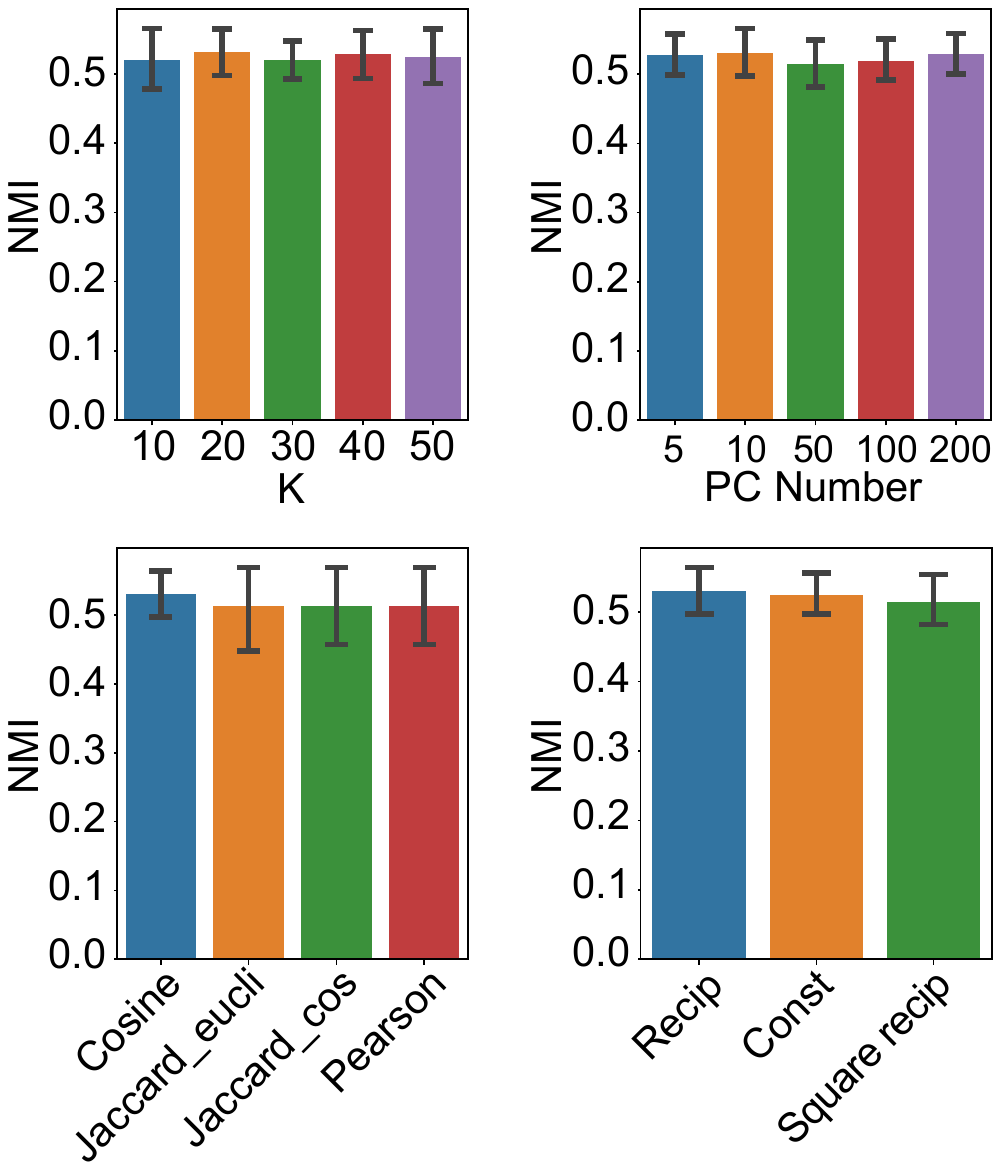

**Supplementary Figure 5.** Using 10,000 SV peaks identified by Descartes and its variants for clustering, we quantitively assessed the impact of different parameters with NMI scores as the metric. The analysis primarily focused on the following four aspects concerning their impact on Descartes: (i) the number of neighboring cells (or spots) considered when constructing the graph of chromatin accessibility (**a**); (ii) the number of principal components (PCs) retained after PCA when constructing the graph of chromatin accessibility (**b**); (iii) the strategies of calculating similarity between neighboring points and the weighting of edges when constructing the graph of chromatin accessibility (**c**); (iv) the relationship between edge weights and cell-cell distance when constructing the graph of spatial locations (**d**).

**Supplementary Figure 6**

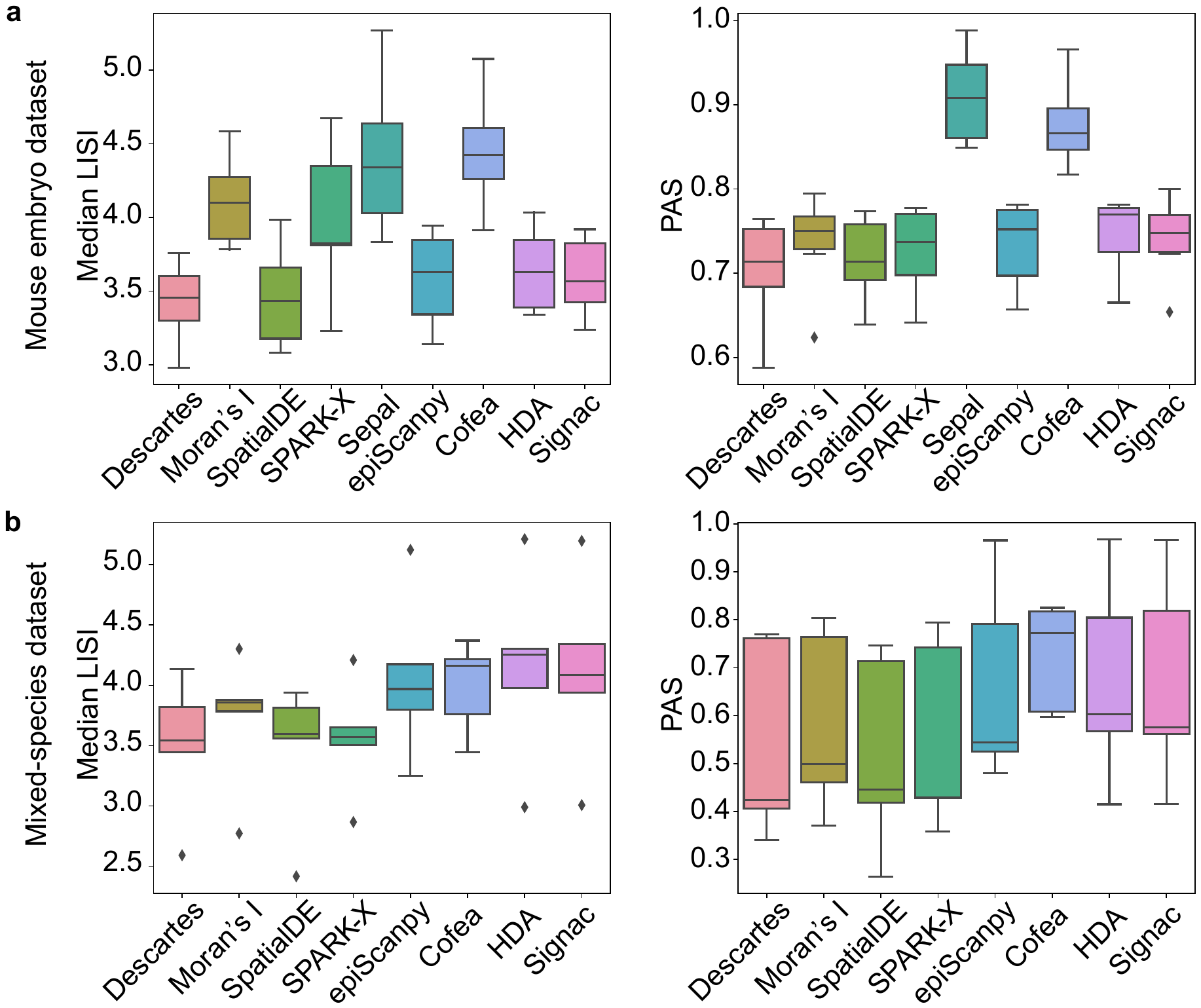

**Supplementary Figure 6.** Clustering performance evaluated by median LISI and PAS scores using SV peaks identified by different methods on the mouse embryo dataset (**a**) and the mixed-species dataset (**b**), respectively.

**Supplementary Figure 7**

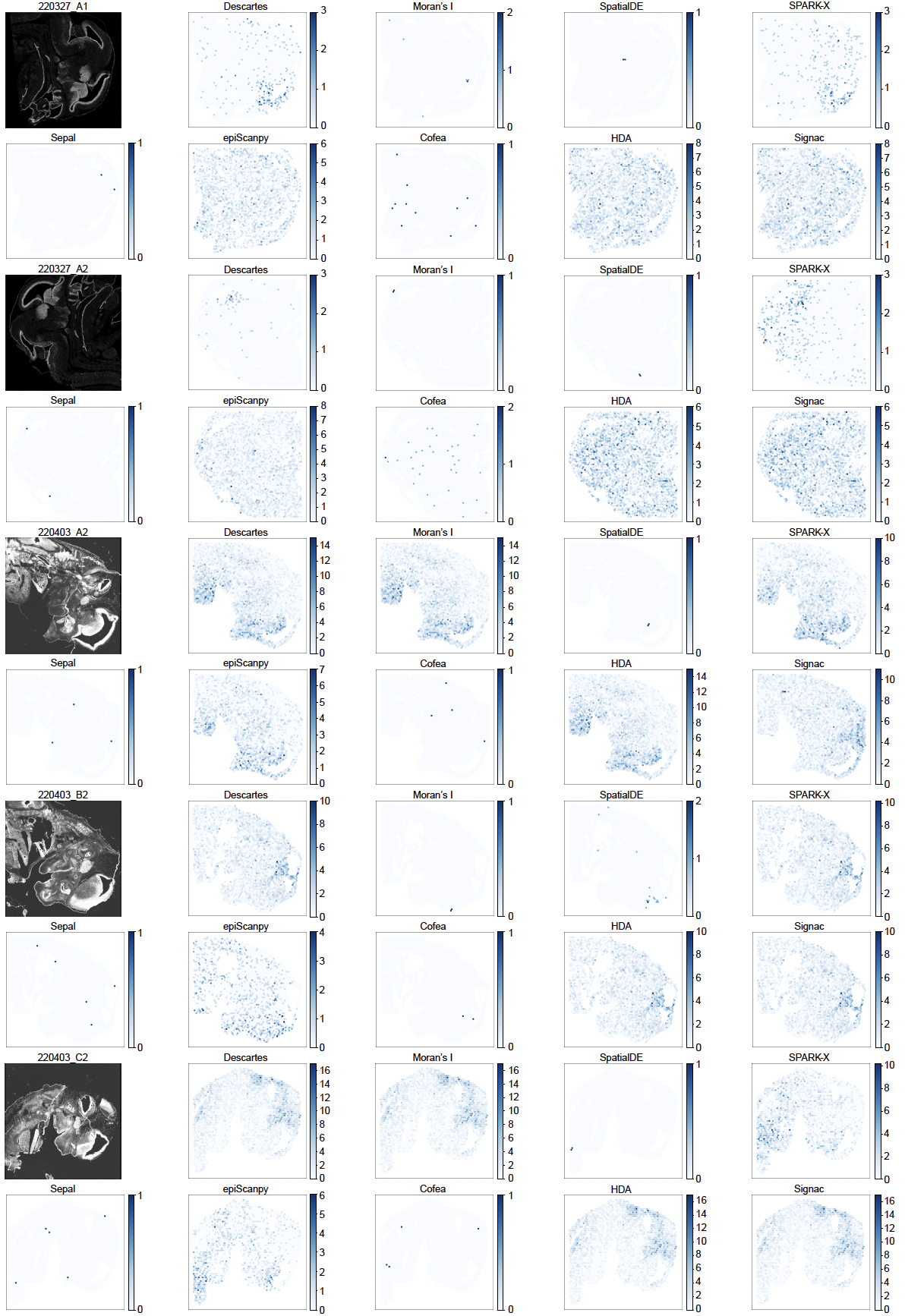

**Supplementary Figure 7.** The top-ranked SV peak identified by each method in the tissue space on the slices from the mouse embryo dataset, compared to histological image (the first subplot).

**Supplementary Figure 8**

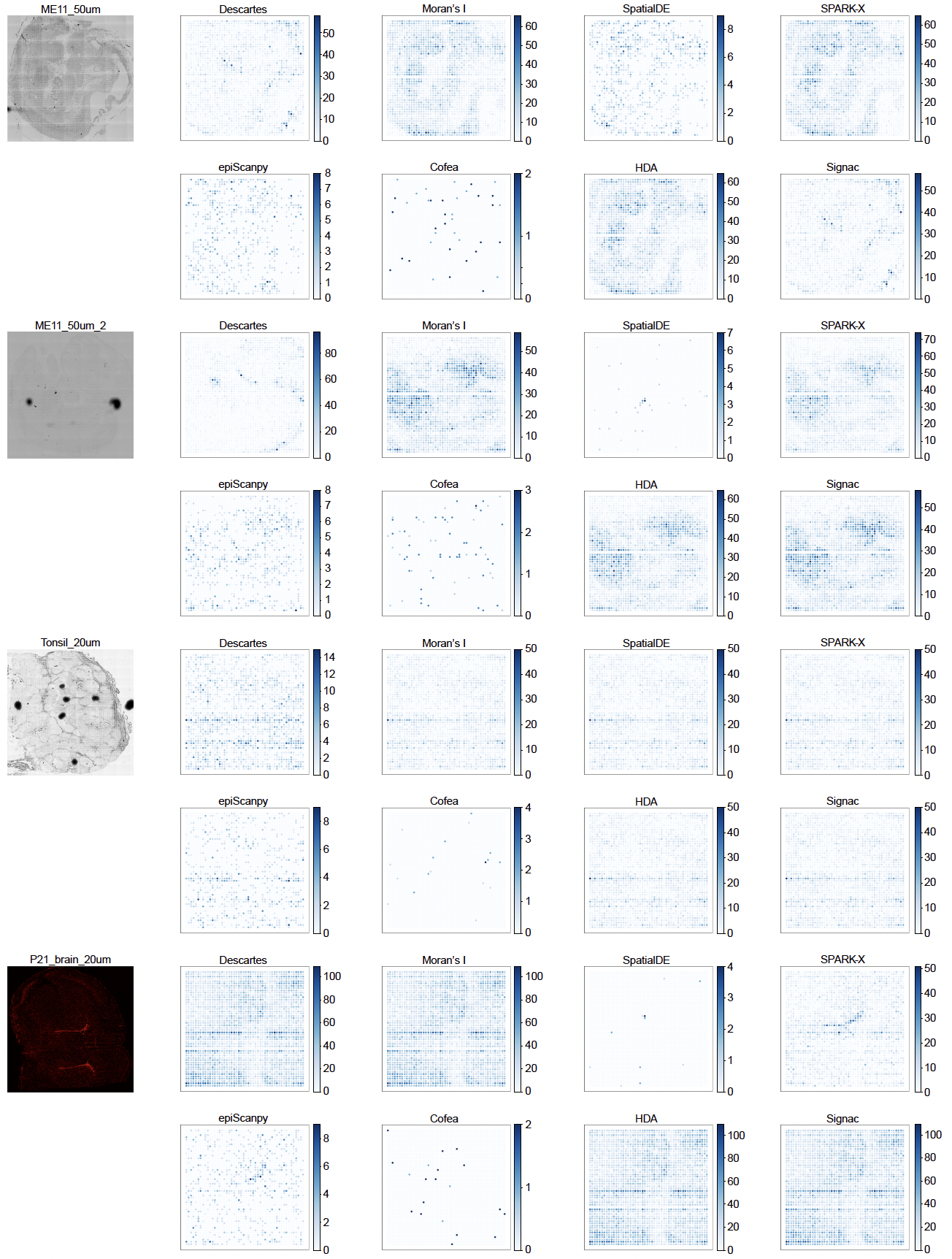

**Supplementary Figure 8.** The top-ranked SV peak identified by each method in the tissue space on the slices from the mixed-species dataset, compared to histological image (the first subplot).

**Supplementary Figure 9**

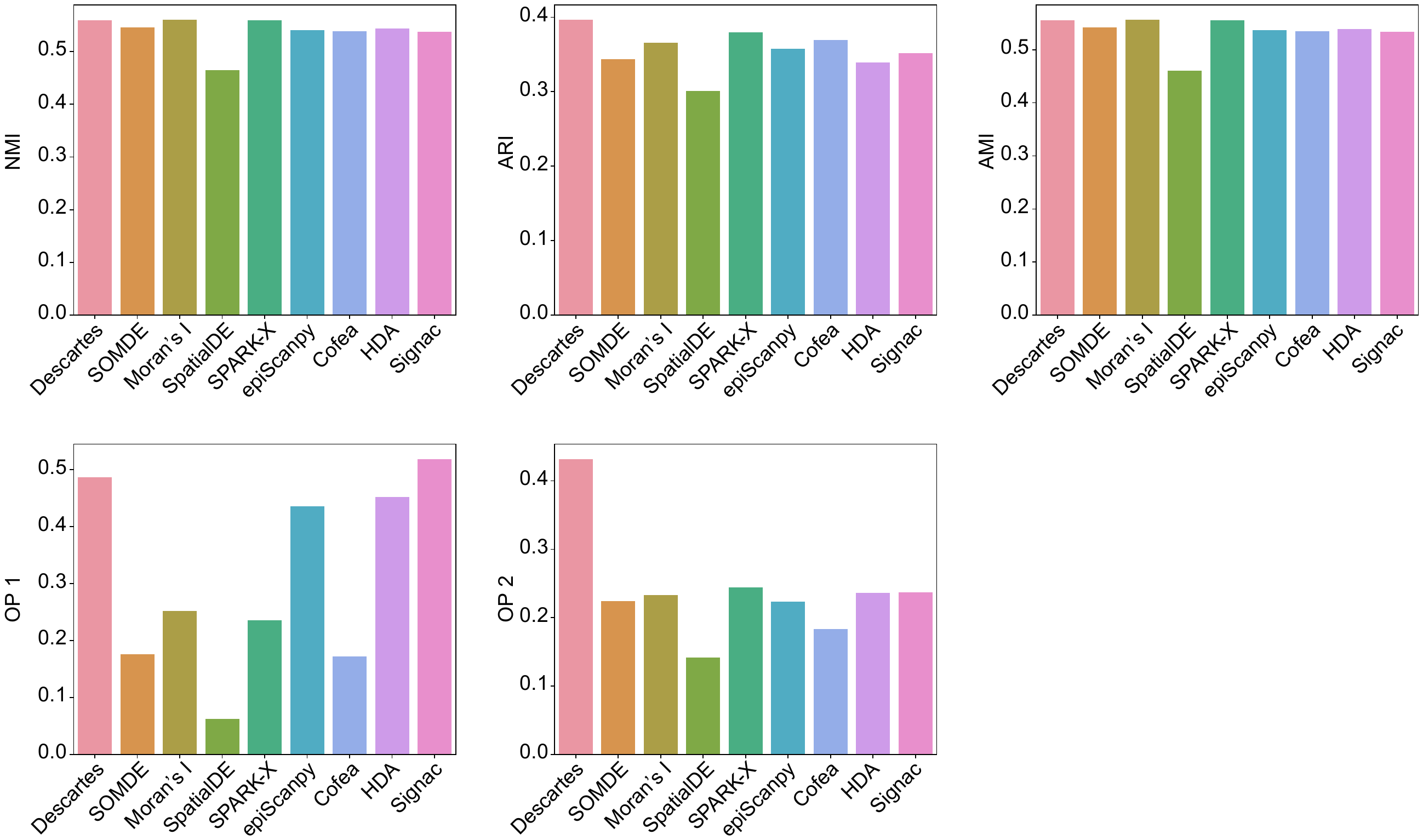

**Supplementary Figure 9.** Benchmarking results before metric transforming of different methods on the metastatic melanoma dataset from two perspectives, that is the ability to facilitate clustering performance and capture domain-specific signals. For the perspective of facilitating clustering performance, we use NMI, ARI and AMI scores as metrics. For the perspective of capturing domain-specific signals, we use OP1 and OP2 scores as metrics.

**Supplementary Figure 10**

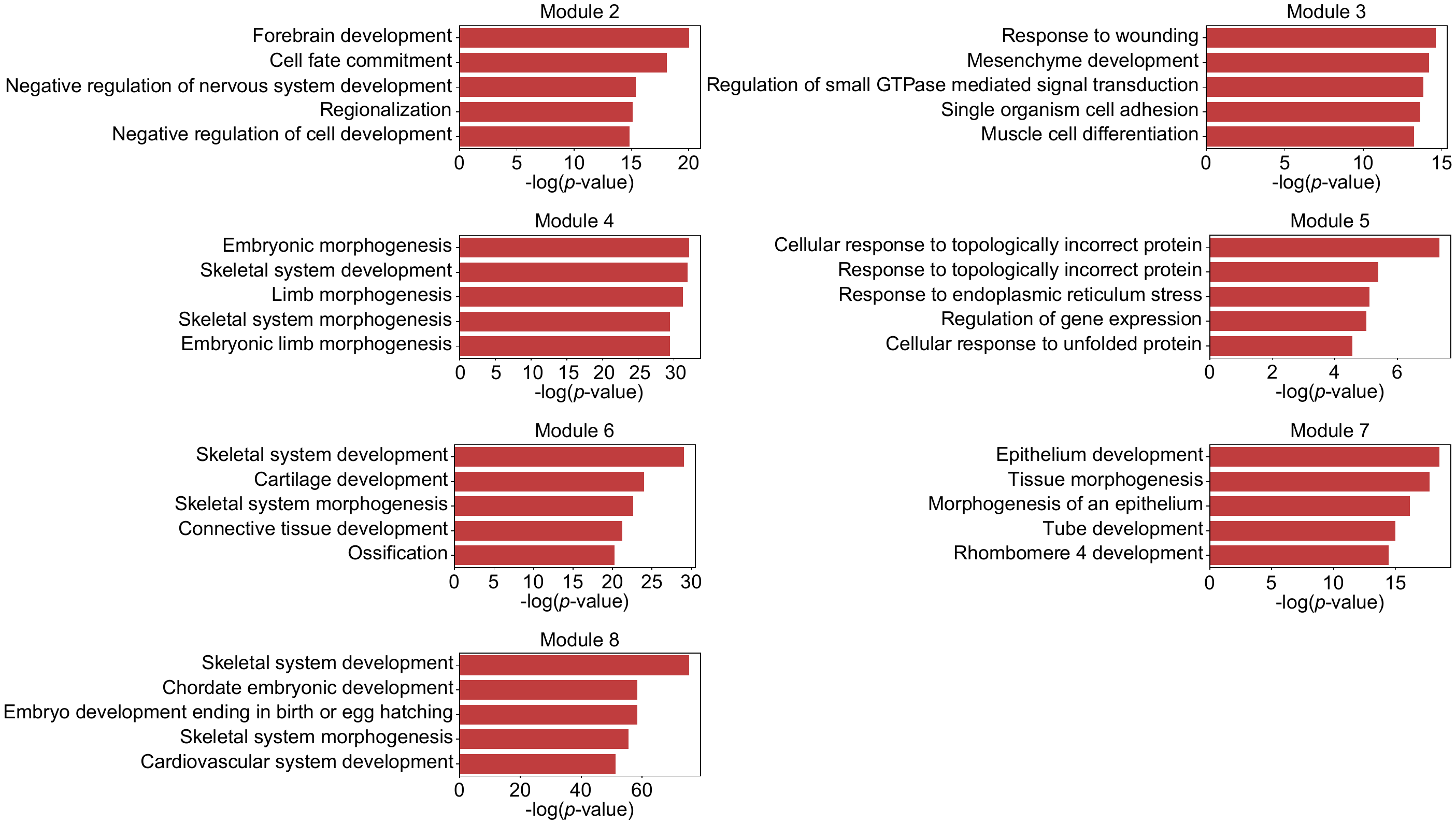

**Supplementary Figure 10.** The top-5 most significant pathways for each module with *p*-values calculated through GREAT analysis. Due to the low number of peaks in peak module 1, there is no significant pathway (*p*-value<0.05) associated with the module.

**Supplementary Tables**

**Supplementary Table 1**

| Dataset | Slices | Spots | Genes | Peaks/Bins | Number of domains | Tissue | Sequencing Technology | Transcriptome data sparsity | Chromatin accessibility data sparsity | Resolution |
| --- | --- | --- | --- | --- | --- | --- | --- | --- | --- | --- |
| Mouse brain dataset | E11_0-S1 | 1258 | 32285 | 242024 | 8 | Brain_E11_0 | MISAR-seq | 0.9252 | 0.9665 | 50 um |
|  | E13_5-S1 | 1777 | 32285 | 271126 | 12 | Brain_E13_5 | MISAR-seq | 0.9106 | 0.9662 | 50 um |
|  | E15_5-S1 | 1949 | 32285 | 265014 | 12 | Brain_E15_5 | MISAR-seq | 0.8947 | 0.9476 | 50 um |
|  | E18_5-S1 | 2129 | 32285 | 294734 | 14 | Brain_E18_5 | MISAR-seq | 0.9343 | 0.9618 | 50 um |
| Mouse embryo dataset | 220327_A1 | 3274 | - | 248047 | - | Embryo_E15.5 | Spatial ATAC | - | 0.9918 | 55 um |
|  | 220327_A2 | 3903 | - | 248047 | - | Embryo_E15.5 | Spatial ATAC | - | 0.9929 | 55 um |
|  | 220403_A2 | 3160 | - | 248047 | - | Embryo_E13.5 | Spatial ATAC | - | 0.9973 | 55 um |
|  | 220403_B2 | 2970 | - | 248047 | - | Embryo_E13.5 | Spatial ATAC | - | 0.9958 | 55 um |
|  | 220403_C2 | 2246 | - | 248047 | - | Embryo_E12.5 | Spatial ATAC | - | 0.993 | 55 um |
|  | 220403_D2 | 2234 | - | 248047 | - | Embryo_E12.5 | Spatial ATAC | - | 0.9947 | 55 um |
| Metastatic melanoma dataset | adata_annotated | 2535 | 36601 | 53431 | 10 | metastatic melanoma | Slide-tags+scRNA-seq+scATAC-seq | 0.9148 | 0.9811 | single cell (less than 10 um) |
| Mixed species dataset | ME11 | 2500 | - | 290412 | - | E11 mouse embryo | Spatial-ATAC-seq | - | 0.9761 | 50 um |
|  | ME13 | 2500 | - | 289542 | - | E13 mouse embryo | Spatial-ATAC-seq | - | 0.9765 | 50 um |
|  | Tonsil | 2500 | - | 248909 | - | human tonsil | Spatial-ATAC-seq | - | 0.9839 | 20 um |
|  | ME11 | 2500 | - | 283043 | - | E11 mouse embryo | Spatial-ATAC-seq | - | 0.9761 | 20 um |
|  | P21_brain | 2500 | - | 258604 | - | mouse postnatal day 21 (P21) brain | spatial ATAC-seq | - | 0.9879 | 20 um |
